# *TOMM40* suppression promotes neuronal cholesterol imbalance and molecular and behavioral phenotypes of Alzheimer’s disease

**DOI:** 10.1101/2025.10.31.685963

**Authors:** Neil V Yang, Shaowei Wang, Boyang Li, Jeffrey Simms, Linsey Dinh, Annie Huang, Jacob H Oei, Hussein N Yassine, Ronald M Krauss

## Abstract

**INTRODUCTION:** While the *APOE4* allele is a major risk factor for Alzheimer’s disease (AD), the role of *TOMM40*—an adjacent gene involved in mitochondrial protein import—is not known.

**METHODS:** Mice, human iPSC-derived neurons (iNeurons), and human brain tissue were used for study of animal cognition, cholesterol metabolism, mitochondrial function, and gene expression.

**RESULTS:** *TOMM40* knockdown (KD) impaired memory in mice and increased cholesterol and Aβ 42 in mouse brains and human iNeurons. KD disrupted mitochondria-endoplasmic reticulum contact sites (MERCs), causing mitochondrial dysfunction and promoting reactive oxygen species that led to activation of LXRB (NR1H2), upregulation of *APOE* and *LDLR.* and increased cellular cholesterol and Aβ 42 independent of *APOE4*. Human brain transcriptomics showed reduced *TOMM40* expression that correlated with cholesterol regulatory gene expression, amyloid burden, and clinical AD diagnosis.

**DISCUSSION:** TOMM40 is a novel mediator of AD pathology through dual effects on MERCs that regulate cholesterol homeostasis and mitochondrial function.

**GRAPHICAL ABSTRACT:** 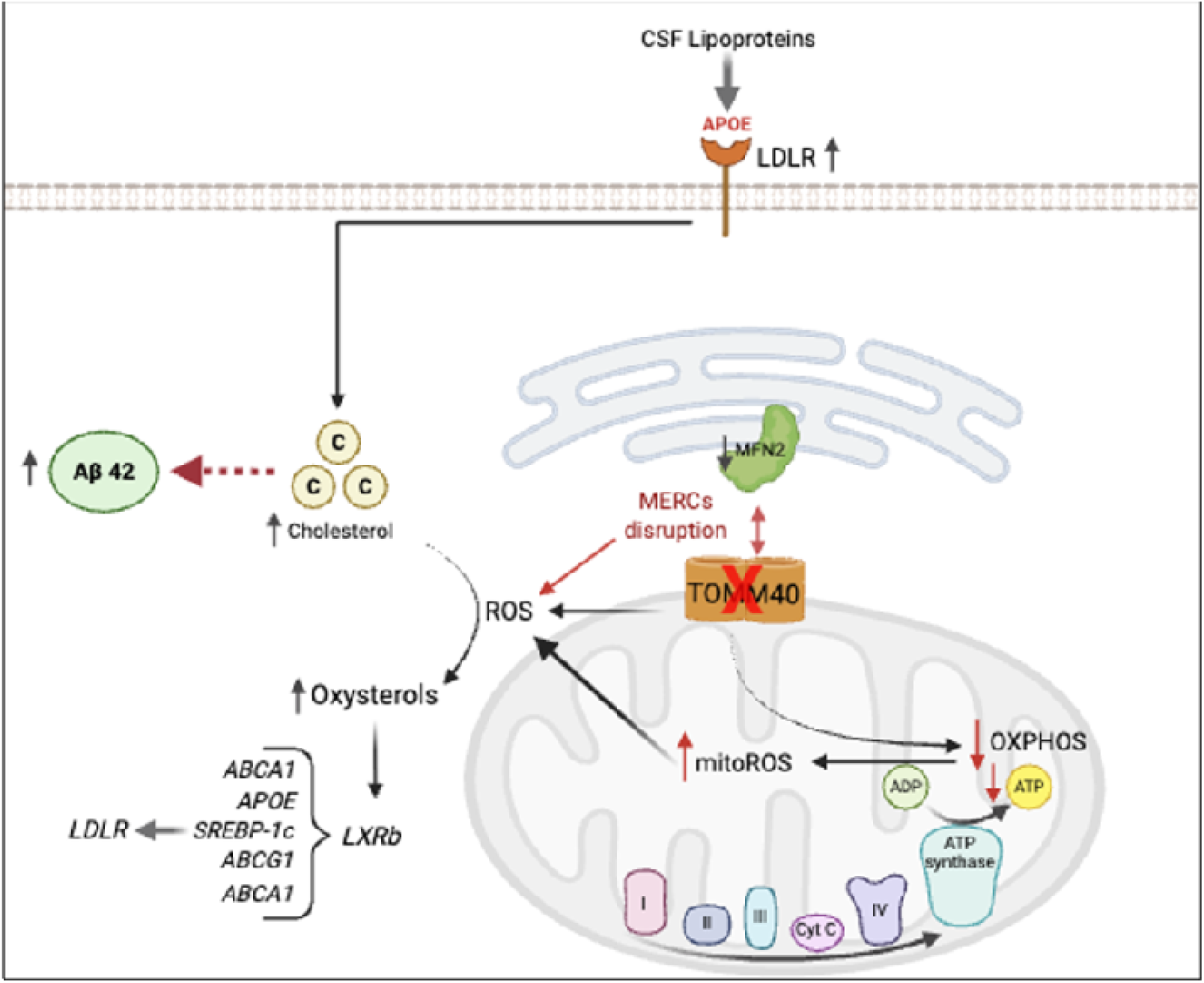

## BACKGROUND

Alzheimer’s disease (AD) is the most common form of dementia, with a worldwide prevalence of 10% in those above age 65^1^. However, the underlying pathologies that lead to development of AD remain unclear. *APOE4*, which encodes an isoform of APOE (apolipoprotein E), is the strongest genetic risk factor for late-onset AD^2^. Approximately 25% of the world population carries an *APOE4* allele, increasing AD risk by 3-4 fold compared to carriers of the *APOE3* “non-pathogenic” allele^3^. APOE is a ligand for all major lipoprotein receptors in the brain including LDLR (low-density lipoprotein receptor), thus promoting uptake of cholesterol from lipoproteins including high-density lipoprotein (HDL) particles, the main transporters of cholesterol in the brain^4^. Notably, there is recent evidence that high brain cholesterol level, quantified by positron emission tomography (PET) scans, is a risk factor for AD, and that brain cholesterol imbalance influences Aß aggregation, phosphorylation of tau proteins, and synaptic dysfunction – all major hallmarks of AD^5,6^.

The gene encoding TOMM40 (translocase of outer mitochondrial membrane 40) is adjacent to *APOE*. Multiple studies have shown that single-nucleotide polymorphisms (SNPs) within *TOMM40* and *APOE* are in strong linkage disequilibrium, and that subsets of these SNPs, most notably the *TOMM40* rs 0524523 polymorphism, have additive associations with late onset AD, independent of *APOE* genotype^7–9^. TOMM40 is the main channel-forming subunit of the translocase of the outer mitochondrial membrane (TOM) complex that is required for the transport of precursor proteins into the mitochondria to maintain mitochondrial function^10^. In this regard, it is notable that mitochondrial dysfunction is present in the brain of AD patients, resulting in increased production of cellular reactive oxygen species (ROS) and subsequent neuronal damage and degeneration^11^. However, despite the associations of *TOMM40* SNPs with AD, and the role of TOMM40 in maintaining mitochondrial function, no studies have yet directly tested whether TOMM40 has a role, either independently or in conjunction with APOE4, in the development of AD.

We have recently shown that hepatic *TOMM40* knockdown (KD) disrupts mitochondria-endoplasmic reticulum contact sites (MERCs), leading to increased reactive oxygen species and oxysterol production resulting in activation of LXR, and increased uptake of cholesterol due in part to LXR-mediated *APOE* upregulation^12^. These findings have raised the possibility that similar effects may result from *TOMM40* KD in neural cells, thus contributing to AD pathophysiology via effects on both cholesterol homeostasis and mitochondrial function, The present study was aimed at testing this hypothesis in human induced pluripotent stem cells (iPSC)-derived neurons and *Tomm40* KD mice.

## METHODS

### Animals

#### Tissue collection and analysis

Eight-week old C57BL/6J male mice (n=6 per group) were obtained from Jackson Laboratory (stock 000664) and placed on a chow-fed diet. At 10 weeks of age, mice received intraperitoneal (IP) injections of either AAV8-*Tomm40* shRNA (4 × 10¹¹ GC) or control AAV8-CMV-null vector (VectorBuilder). Body weight and food intake were monitored weekly. At 16 weeks, mice were euthanized in the unfasted state, and brain samples were collected, snap-frozen in liquid nitrogen, and stored at −80 °C. For electron microscopy, tissues were fixed immediately postmortem in 2% glutaraldehyde and 2% paraformaldehyde. All animal studies were reviewed and approved by the Institutional Animal Care and Use Committee (IACUC) at the University of California, San Francisco (approval number: AN196014-01B).

#### Behavioral experiments

Forty 8-wk old male C57B6/J mice (Jackson Laboratory, stock 000664) were group housed in cages of five mice, exposed to a 12-hr light-dark cycle, and kept in a humidity and temperature-controlled environment with *ad libitum* access to food and water. At 10 wks of age, mice were injected with either *Tomm40* (VectorBuilder Inc., Chicago, IL) or control AAV. Behavioral assessments were initiated 6 wks following the delivery of the plasmids. The mice were tested in two cohorts of 20 mice. The first cohort was assessed in the Elevated Plus Maze, Open Field, Novel Object Recognition with a 24 hr delay, and the Object Context Congruence Task with a 4 hr delay. The second cohort was tested using the same behavioral test battery except that the OCC was replaced with the Morris Water Maze.

All procedures were conducted in a randomized and blinded manner. Animal experiments were approved by the University of California, San Francisco, Laboratory Animal Resource Center.

#### Elevated Plus Maze

The Elevated Plus Maze (EPM) apparatus (Kinder Scientific, Chula Vista, CA) is 63 cm above the ground, composed of two open arms (without walls, 36 cm long x 6 cm wide), two closed arms (36 cm long x 6 cm wide with 16 cm high walls), with the intersection being 6 cm x 6 cm wide. Before testing, the mice were habituated in the room under dim lighting for an hour. After habituation, the mice were tested by placing them in the intersection of the open and closed arms to explore the maze for 10 mins. Data was recorded using Kinder Scientific Motor Monitor software. Total distance and the time spent in the open arms were monitored with infrared photobeams.

#### Open Field Activity

Spontaneous locomotor activity was accessed using the Med Associates Seamless Open Field (OF) System and Activity Monitor Software (Med Associates, Fairfax, VT). The apparatus consists of a clear acrylic chamber (43 x 43 x 30 cm) that is housed in a larger sound and light attenuating shell to block out external stimuli. Within the clear acrylic chamber, horizontal and vertical movement is detected automatically through two 16 x 16 photobeam arrays. Mice were habituated for an hour before testing under normal light. During testing, mice were placed in the center of the chamber and were able to freely explore for 15 mins. Total movement, number of rearings, ratio of movements in center movement versus the periphery, and resting by zone were recorded.

#### Novel Object Recognition

Novel Object Recognition (NOR) was utilized to measure visual recognition and declarative memory. The test was conducted in 30 x 30 cm grey acrylic chambers with two objects placed in the center of the box, equidistant from each other and the walls of the chamber. The objects used in this experiment were two distinct 6-8 cm LEGO DUPLO towers with 20 µl of undiluted vanilla on cotton swabs inside each object to increase exploration. Object pairs were counterbalanced throughout the cohort of mice. On the first day, the mice were brought into the testing room to habituate under normal lighting conditions for an hour. For the training trial, the mice were put in the center of the chamber with two identical objects, where they freely explored the chamber and interacted with the objects for 10 mins. Twenty-four hrs after the training trial, the mice were brought back into the same room to habituate again for an hour under normal lighting. In the testing session, the mice were placed into the center of the same chamber to interact and explore for another 10 mins, with one of the two objects replaced with a novel object. All training and testing trials were video recorded with EthoVisionXT video tracking system (Noldus, Wageningen, Netherlands), then analyzed using CleverSys TopScan (Reston, VA) where the number of sniffing bouts and duration of sniffing were recorded.

#### Object Context Congruence Task

The Object Context Congruence (OCC) Task is a pattern separation/pattern completion task. Briefly, mice were moved into the testing room for 1 hr under normal lighting conditions to habituate. The task was conducted inside sound attenuating chambers equipped with two 5 W houselights and an overhead video camera. Mice were randomly assigned to two groups: Context 1 first or Context 2 first for trial one, then the opposite for trial two. The contexts are square open top chambers (30 cm x 30 cm x 25 cm) made of smooth gray acrylic. Objects were placed side by side in the center of the arena. Context 1 consisted of a plain white chamber, two identical objects, X_1_ and X_2_ (white rectangular blocks with tented tops), and was cleaned with 70% ethanol. Context 2 consisted of a chamber with checkered walls, two identical objects, Y_1_ and Y_2_ (50mL Erlenmeyer flask filled with colored paper) and was cleaned with 1% Acetic Acid. The mice were tested within one day with three 10-min trials. In the first trial the mice explored either Context 1 or Context 2 for 10 mins, then were returned to their home-cage for 30 mins. For the second trial, the mice explored the context they were not exposed to during the first trial for 10 mins, then were returned back to their home-cage for 4 hrs. Following the four delay, the mice were put back into the trial two context for the 10-min test trial. During the test phase, the incongruent objects were the X object placed in Context 2 and the Y object placed in Context 1. Each trial was video recorded from above using EthoVisionXT video tracking system (Noldus, Wageningen, Netherlands) and analyzed with CleverSys TopScan (Reston, VA) to record interaction bouts and duration of sniffing on each of the objects.

#### Morris Water Maze

To assess spatial learning and memory in mice treated with the *Tomm40* shRNA AAV, we performed the Morris Water Maze. The water maze apparatus consists of a pool (122 cm diameter; 50 cm high), filled with water maintained at 22 ± 1°C, made opaque with non-toxic white tempera liquid paint. On the four walls of the room are distinct, large black and white extramaze cues. The same person completes the entire training paradigm, avoiding heavily scented soaps, lotions, or perfumes and avoids changing such items. Water Maze procedure is broken up into four different portions over the course of 10-12 days: pre-training, hidden platform training, probe trials, and cued-platform trials. All trials were video recorded with an EthoVisionXT video tracking system (Noldus, Netherlands).

#### Pretraining

Before hidden platform training began, mice were given four consecutive pre-training trails. During these trials, the mice swam down a rectangular training track (15 x 122 cm) onto a ramp connected to a platform (15 x 15 cm), that is 1.5 cm below the water at the end of the track. The mice had 60 secs to find the platform; if they did not find the platform within 60 secs, they were then guided to the platform by the experimenter’s hand. Mice were dried with a paper towel and single housed for the duration of the experiment. If a mouse did not groom within 3 mins after returning to a cage, a heat lamp is provided to prevent hypothermia.

#### Hidden Platform Training

Hidden platform training spans across five consecutive days, with two sessions of two trials each day. Between these sessions there was a 2 to 3 hr inter session interval (ISI) and between the trials there was a 15–20 min inter-trial interval (ITI). The mouse was dropped into the pool facing the wall. During the trials the mouse was given 60 secs to find a hidden platform. The platform location remained the same over the five days and was submerged 1.5 cm under the water without any local visible cues. The location of where the mouse was dropped was randomized across trials, but each mouse was dropped at a consistent location within the trial. The drop locations were randomized so that each two-trial session contains a drop location that is “closer” and “farther” from the platform. Once the mouse found and remained on the platform, the trial was stopped and the mouse remained on the platform for 10-15 secs before returning to its home cage. If the mouse left the platform before 10-15 secs or if 60 secs had elapsed without the mouse finding the platform, the mouse was guided on to the platform with the experimenter’s hand until it remained there for 10-15 secs. Mice were dried with a paper towel and returned to their home cage. A heat lamp/pad was provided to mice that did not groom within 3 mins to prevent hypothermia.

#### Probe

The probe trials were performed 24, 72, and 120 hrs after the final day of hidden platform training. The hidden platform was removed and the mouse was dropped facing the wall, 180° from where the hidden platform used to be. The mouse’s swim pattern was recorded for a single 60 sec trial. After 60 secs, the experimenter placed their hand where the platform was and let the mouse swim to their hand. The same procedures were used to dry the mouse and prevent hypothermia as mentioned previously.

#### Cued Platform Trials

Cued platform trials were performed after the final day of probe. There were three sessions of two trials each, with a 1.5 hr break between each session and a 10-15 min break between each trial. The cued platform was submerged 1.5 cm under the water, while a black and green stripped mast (15 cm in height) was used as a visual cue for the platform. The platform was moved to a different quadrant of the pool for each two-trial session. These quadrants were the three quadrants not used for hidden platform training. The drop locations varied between each set of two trials, corresponding to which quadrant the platform was in. All mice received the same cued platform location and two drop locations for the session. The mice were dropped facing the wall and were given 60 secs to find the visible platform and remain on it for 10-15 secs. If the mouse failed to find the platform, they were guided to the platform by the experimenter’s hand and remained on the platform for 10-15 secs.

### iPSCs Maintenance & Differentiation

Human derived induced-pluripotent stem cells (iPSCs) were acquired from the Jackson Laboratories (JAX). The APOE3 homozygous (control) and APOE4 homozygous isogenic cell lines were derived from KOLF2.1J reference cell line and are commercially available. All iPSC cell lines were induced into neural progenitor cells (NPCs) according to the protocol of Bardy et al^13^. NPCs were plated at a density of 1.5-2 × 10^5^ cells/mL in 6-well plates coated with poly-L-ornithine/laminin and incubated in supplemented STEMdiff™ Neural Progenitor Basal Medium (STEMCELL Technologies). Once the NPCs were ∼80% confluent, NPCs were differentiated using STEMdiff™ Brain Phys Media supplemented with BDNF, GDNF, B27, Ascorbic Acid, laminin, and cAMP (STEMCELL Technologies), media change occurred daily. The media was changed 2-3 times a week for 3 weeks until the neurons were fully differentiated. Mature neurons were verified by NeuN protein expression using western blot^14^.

### siRNA Transfections

Once iNeurons were fully differentiated and mature, siRNA transfections were used to perform knock-down (KD) of *TOMM04, MFN2, APOE, LDLR,* and *LRP1* in iNeurons by addition of their respective siRNAs (10 µM) using Lipofectamine STEM transfection reagent (Life Technologies) and Opti-MEM I (Gibco) according to the manufacturer’s instructions for 48 h at 37 °C.

### Lipid Extraction and cholesterol quantification

Mouse brain samples were homogenized using a GentleMACS™ dissociator (Miltenyi Biotec), and cells were lysed with a cell disruptor (Bio-Rad) in either a chloroform:methanol:water mixture (8:4:3, v/v/v) for total lipid extraction^15^, or hexane:isopropanol (3:2, v/v) for cholesterol extraction^16^. For cholesterol analysis, samples were dried under nitrogen gas and reconstituted in buffer (0.5 M potassium phosphate, pH 7.4, 0.25 M NaCl, 25 mM cholic acid, 0.5% Triton X-100). Intracellular cholesterol was then quantified using the Amplex Red Cholesterol Assay Kit (Life Technologies), according to the manufacturer’s protocol.

### Enzyme-linked immunosorbent assay (ELISA)

Cells and isolated brain (hippocampus) tissues were lysed in M Cellytic Lysis Buffer supplemented with 1% protease inhibitor (Halt™ Protease Inhibitor Cocktail; ThermoFisher Scientific) for 15 mins using a cell disruptor or homogenizer. Lysates were centrifuged at 14,000 × g for 15 mins, and the supernatant was collected for analysis. Aß 42 and p-tau protein levels were quantified using ELISA kits from Thermofisher Scientific, following the manufacturer’s instructions. All measurements were normalized to total protein content, assessed by BSA assay (Genesee Scientific).

### Mitochondrial bioenergetics

NPCs were seeded at a density of 2,000 cells per well in 96-well plates and underwent differentiation for 2 weeks. Once differentiated, iNeurons were incubated in phenol-free neuron differentiation medium (STEMCELL) supplemented with 2 mM sodium pyruvate (Gibco), 2 mM GlutaMAX™ (Gibco), and 10 mM glucose (Sigma), adjusted to pH 7.4. Mitochondrial function was assessed using the Seahorse XFe96 Extracellular Flux Analyzer (Agilent). During the assay, 1.5 μM oligomycin, 2 μM FCCP, and 2 μM antimycin A + rotenone (Seahorse XF Cell Mito Stress Test Kit, Agilent) were sequentially injected to evaluate basal respiration, maximal respiration, ATP production, and proton leak. Oxygen consumption rate (OCR) values were corrected for non-mitochondrial respiration and normalized to total protein content per well, quantified by BCA assay (Genesee Scientific).

### Mitochondrial assays

Cellular reactive oxygen species (ROS) levels were measured using the DCFDA/H DCFDA Cellular ROS Assay Kit (Abcam), following the manufacturer’s instructions. Cells were incubated with DCFDA for 45 mins, then washed with 1× DPBS. Fluorescence was detected using a spectrophotometer at an excitation/emission wavelength of 485/535 nm.

To detect mitochondria-derived ROS, iNeurons were incubated with 5 µM MitoSOX™ Red (Molecular Probes) for 20 min at 37 °C. Cells were rinsed twice with pre-warmed 1X phosphate-buffered saline (PBS) and was replaced with phenol red-free DMEM (21063029; Gibco) supplemented with glucose and sodium pyruvate. Fluorescence was quantified at 396/610 nm using an Agilent BioTek™ microplate fluorescence spectroscopy reader. All measurements were normalized to total protein content, assessed by BSA assay (Genesee Scientific).

### Transmission electron microscopy

#### Sample preparation

NPCs were incubated on MatTek glass-bottom dishes (P35G-1.5-14-C, MatTek) and differentiated into iNeurons after 2-weeks before being fixed in 2% glutaraldehyde combined with 2% paraformaldehyde (prepared by the Electron Microscopy Lab, UC Berkeley) for 24 hrs. Following fixation, both cells and brain tissues were post-fixed in 1% osmium tetroxide prepared in 0.1 M sodium cacodylate buffer (pH 7.2) for 1–2 hrs. Samples were then dehydrated through a graded ethanol series (30% to 100%). Brain tissues were embedded in increasing concentrations of Durcupan resin. iNeurons were infiltrated with 50% Epon-Araldite resin containing benzyldimethylamine (BDMA) as an accelerator, followed by 100% resin for 1 hr each. All samples were polymerized at 60 °C for 24 hrs.

#### Imaging

Resin-embedded sample blocks were trimmed, and 90 nm ultrathin sections were prepared using a Leica UC6 ultramicrotome (Leica Microsystems, Vienna, Austria). Sections were collected onto Formvar-coated slot grids. Target regions were identified and imaged using a Tecnai 12 transmission electron microscope (120 kV; FEI, Hillsboro, OR, USA). Images were acquired with a Gatan Rio16 CMOS camera and processed using GMS3 software (Gatan Inc., Pleasanton, CA, USA).

#### Image Processing

TEM images were analyzed with ImageJ software according to the method for mitochondrial morphology quantification of Lam et al^17^. Mitochondria-ER contact sites were manually identified from high-resolution, high-magnification TEM images. Structural parameters were quantified by outlining contact regions using an optical pen in ImageJ. Mitochondria–ER contact sites (MERCs), as well as other organelle interfaces, were defined by a 10–30 nm gap between the outer mitochondrial membrane and either the ER or lipid droplet membrane, with MERCs further distinguished by the absence of ribosomes on the adjacent ER membrane^18^.

### Fluorescence imaging

To assess HDL uptake, iNeurons were incubated with 10 μM DiI-labeled HDL (Kalen Biomedical LLC), derived from human plasma, for 4 hrs at 37 °C. Following incubation, cells were washed and resuspended in 1X PBS. Fluorescence was measured using a spectrophotometer and normalized to total protein content per well, determined by BCA assay (Genesee Scientific).

### RT-qPCR

RNA was isolated from brain tissues and cell samples using the RNeasy Mini Qiacube Kit (Qiagen) with the Qiacube Connect system, following the manufacturer’s instructions. Complementary DNA (cDNA) was synthesized from total RNA using the High Capacity cDNA Reverse Transcription Kit (Applied Biosystems). Gene-specific primers, designed and obtained from Elim Biopharmaceuticals, were used with SYBR™ Green qPCR Master Mix (Thermo Fisher Scientific) on an ABI PRISM 7900 Sequence Detection System to quantify mRNA expression levels. Primer sequences used in this study are provided in **Table S1**. The average of triplicate measurements for each sample was normalized to GAPDH (human) or 18s rRNA (mouse) as housekeeping genes.

### Immunoblotting

Cells and brain tissues were lysed in M Cellytic Lysis Buffer supplemented with 1% protease inhibitor (Halt™ Protease Inhibitor Cocktail; ThermoFisher Scientific) for 15 mins using a cell disruptor or homogenizer, respectively. The lysates were then centrifuged at 14,000 × g for 15 mins, and the supernatants were collected. Protein concentrations were determined using a BCA assay (Genesee Scientific). Proteins were separated on 4–20% Tris-polyacrylamide gradient gels (Bio-Rad) and transferred onto nitrocellulose membranes using the iBlot™ 2 Gel Transfer Device (ThermoFisher Scientific). Membranes were blocked in Tris-buffered saline with 0.1% Tween-20 (TBST) containing 5% milk for 2 hrs to reduce nonspecific antibody binding. Subsequently, membranes were incubated overnight at 4 °C on a rotating platform with primary antibodies diluted 1:1000 (v/v) in TBST. Following washes with TBST, membranes were incubated for 30 mins with HRP-conjugated secondary antibodies—anti-rabbit IgG (7074) and anti-mouse IgG (7076) from Cell Signaling Technology—diluted 1:2500 (v/v), then washed again. Primary antibodies used in this study are provided in **Table S2**. Protein bands were visualized using SuperSignal™ West Pico PLUS Chemiluminescent Substrate (ThermoFisher Scientific).

### Bulk-RNA sequencing analysis

Filtered raw count data were obtained from the Synapse AD Knowledge Portal (https://www.synapse.org/#!Synapse:syn9702085) under Synapse ID: syn8456637. The source data were derived from 632 individuals enrolled in the Religious Orders Study and the Rush Memory and Aging Project (ROSMAP)^19^. Both studies received approval from the Institutional Review Board at Rush University Medical Center. All participants provided informed consent, repository consent, and consent under the Anatomical Gift Act. At the time of death, clinical diagnoses classified 83 individuals as having no cognitive impairment (NCI), 162 with mild cognitive impairment (MCI), and 151 with Alzheimer’s disease (AD) dementia^20–22^. CERAD and Braak staging were used to assess neuritic plaque burden and neurofibrillary tangle distribution, respectively^23,24^. APOE genotyping was performed as previously described^25^. The R package GSVA (method option “ssGSEA”) was used to calculate pathway enrichment score of each subject. The annoted gene sets were retrieved from MsigDB. The pathway score was tested to assess differences between the variable of interest.

### Single nucleus-RNA sequencing analysis

Post-quality control (QC) count data and ROSMAP metadata were downloaded from the Synapse AD Knowledge Portal (https://www.synapse.org/#!Synapse:syn52293417) under Synapse ID: syn2580853^26^. The dataset includes samples from 427 individuals enrolled in the ROSMAP cohort, all derived from the dorsolateral prefrontal cortex (DLPFC) of post mortem human brain samples^27^. Nuclei were isolated from frozen postmortem brain tissue and analyzed using droplet-based single-nucleus RNA sequencing (snRNA-seq). The R package GSVA (method option “ssGSEA”) was used to calculate pathway enrichment score of each subject. Singe sample GSEA (ssGSEA) is a non-parametric method to assess normalized difference in empirical cumulative distribution functions (CDFs) of gene expression ranks inside and outside the gene set, representing the enrichment score for the gene set. The annoted gene sets were retrieved from MsigDB.

### Statistical Analysis

All data are expressed as mean ± standard error of the mean (SEM). The N-values shown in the figures represent biological replicates, with a minimum of three replicates performed per condition and experiment. For comparisons between two groups, statistical significance was assessed using Student’s t-test. For analyses involving more than two groups, one-way ANOVA or two-way ANOVA followed by Tukey’s post hoc test was applied. All statistical analyses were conducted using GraphPad Prism 9 (GraphPad Software, Inc.). A p-value less than 0.05 was considered statistically significant.

## RESULTS

### Phenotypes of Alzheimer’s disease pathogenesis in the brains of *Tomm40* KD mice

To assess whether *in vivo Tomm40* suppression induces phenotypic traits of AD, we injected AAV8-*Tomm40* shRNA lentivirus into 10-week old C57BL/6J male mice, resulting in ∼60% reduction of *Tomm40* expression in brain as well as liver, skeletal muscle, and white adipose tissue, after 6-wks post-injection (**Fig. 1A**). Brain cholesterol, particularly cholesterol ester, was significantly increased compared to the control AAV (**Fig. 1B-D**). Furthermore, in these mice we found that *Tomm40* KD had increased Aß 42 protein levels in the brain (**Fig. 1E**). However, there was no change in p-tau protein levels (**Fig. 1F**), indicating either that Tomm40 may not impact p-tau in this transient mouse KD model or that KD efficiency was not sufficient to induce an effect.

**Figure 1.**
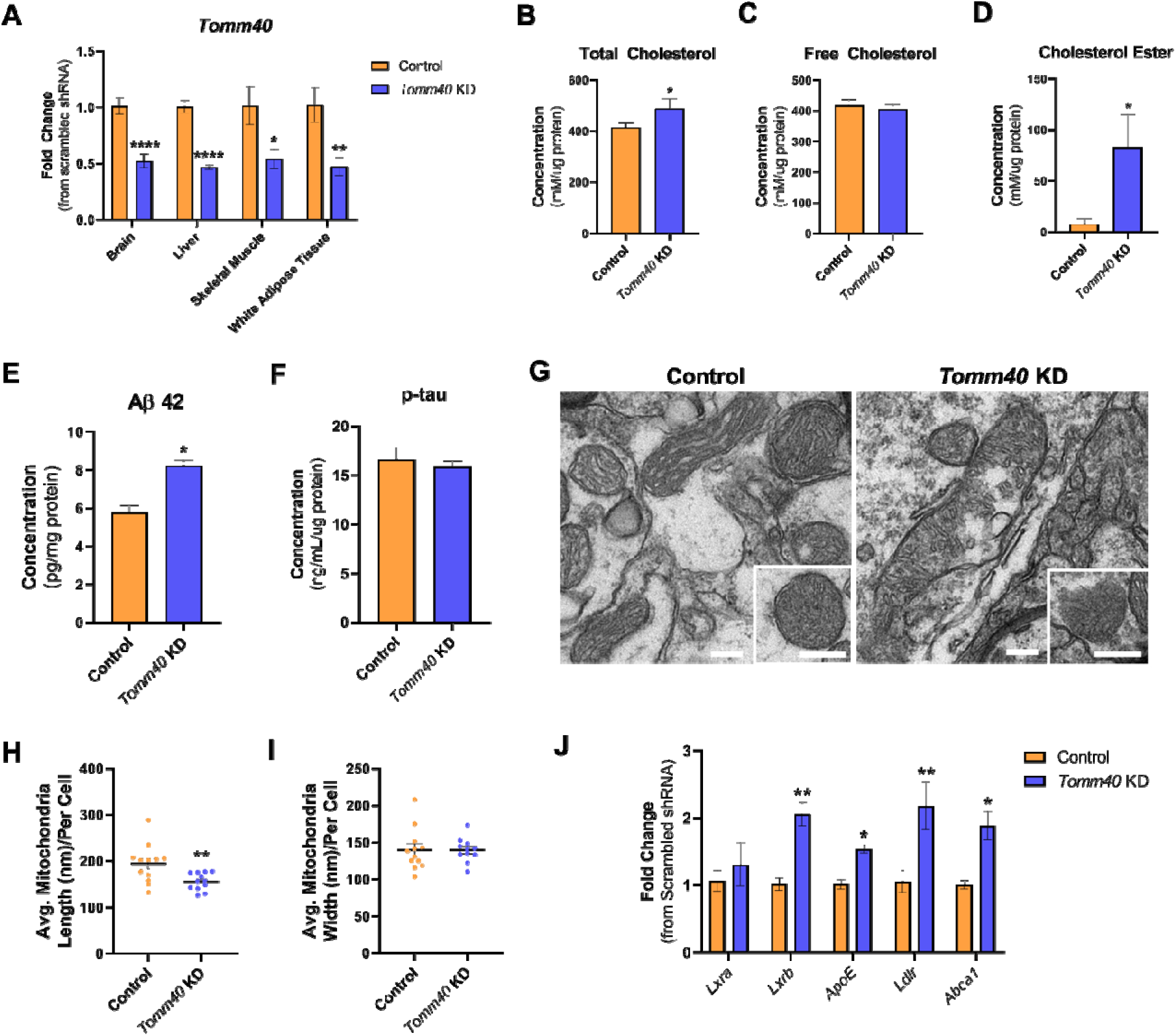
AAV8-*Tomm40* shRNA injected C57BL/6J mice show increased brain cholesterol and A 42 levels, and impaired mitochondrial morphology. (A) Relative mRNA transcript levels of *Tomm40* confirming KD in brain, liver, skeletal muscle, and white adipose tissue of male mice at ∼60% efficiency, quantified by qPCR. Brain total cholesterol (B), free cholesterol (C), and cholesterol ester (D) levels quantified from male mice brain, using Amplex Red Cholesterol Assay. A 42 (E) and phosphorylated-tau (F) levels quantified from male mice brain by enzyme-linked immunosorbent assay. (G) TEM micrographs (scale bar: 250nm) of control vs. *Tomm40* KD AAV injected mice using Image J software: (G) mitochondrial length (nm) and (H) mitochondrial width (nm). (*n=8-12* cells) For all: *n=6* mice/group. *\*p < 0.05, **p<0.01, ****p<0.0001* vs. control AAV by one-way ANOVA, with post-hoc Student’s t-test to identify differences between groups. Data are represented as mean SEM.

Transmission electron microscopy (TEM) of brain tissue in *Tomm40* KD mice revealed damaged and significantly smaller mitochondria (**Fig. 1G-I**), known hallmarks of AD. Moreover, as we previously showed in liver, *Tomm40* KD in male mice promoted brain expression of *Lxrb* and its downstream transcriptional targets including *Apoe*, *Ldlr* regulator of cellular cholesterol uptake, and *Abca1* (ATP binding cassette subfamily A member 1) regulator of expression of genes mediating cellular cholesterol efflux (**Fig. 1J**)^28^.

### Behavioral and cognitive testing of *Tomm40 KD* mice

#### Tomm40 KD results in anxiety-like behaviors in the elevated plus-maze

Testing was performed in 16-week old adult male mice following KD of *Tomm40* using shRNA AAV as described above. With open field testing, there were no differences vs. control AAV in total movement (**Fig. S1A**), center-to-total movement ratio (**Fig. S1B**), rearing (**Fig. S1C**), or resting duration (**Fig. S1D**), suggesting no differences in general locomotor ability^29^. In the EPM, which assesses anxiety-like behaviors^30^, though no differences were found in total distance travelled in the maze (**Fig. 2A**), there was a significant decrease in distance traveled (**Fig. 2B**) and reduction in arm-to-total distance ratio in the open arms of the maze (**Fig. 2C**), indicating an anxiogenic response to the apparatus^31,32^. Finally, while there were no differences in number of open arm entries of the maze (**Fig. 2D**), *Tomm40* KD mice showed increased movement (**Fig. 2E & F**) and entries into the closed arms of the maze (**Fig. 2G)**, indicating increased anxiety-like behaviors^33^.

**Figure 2.**
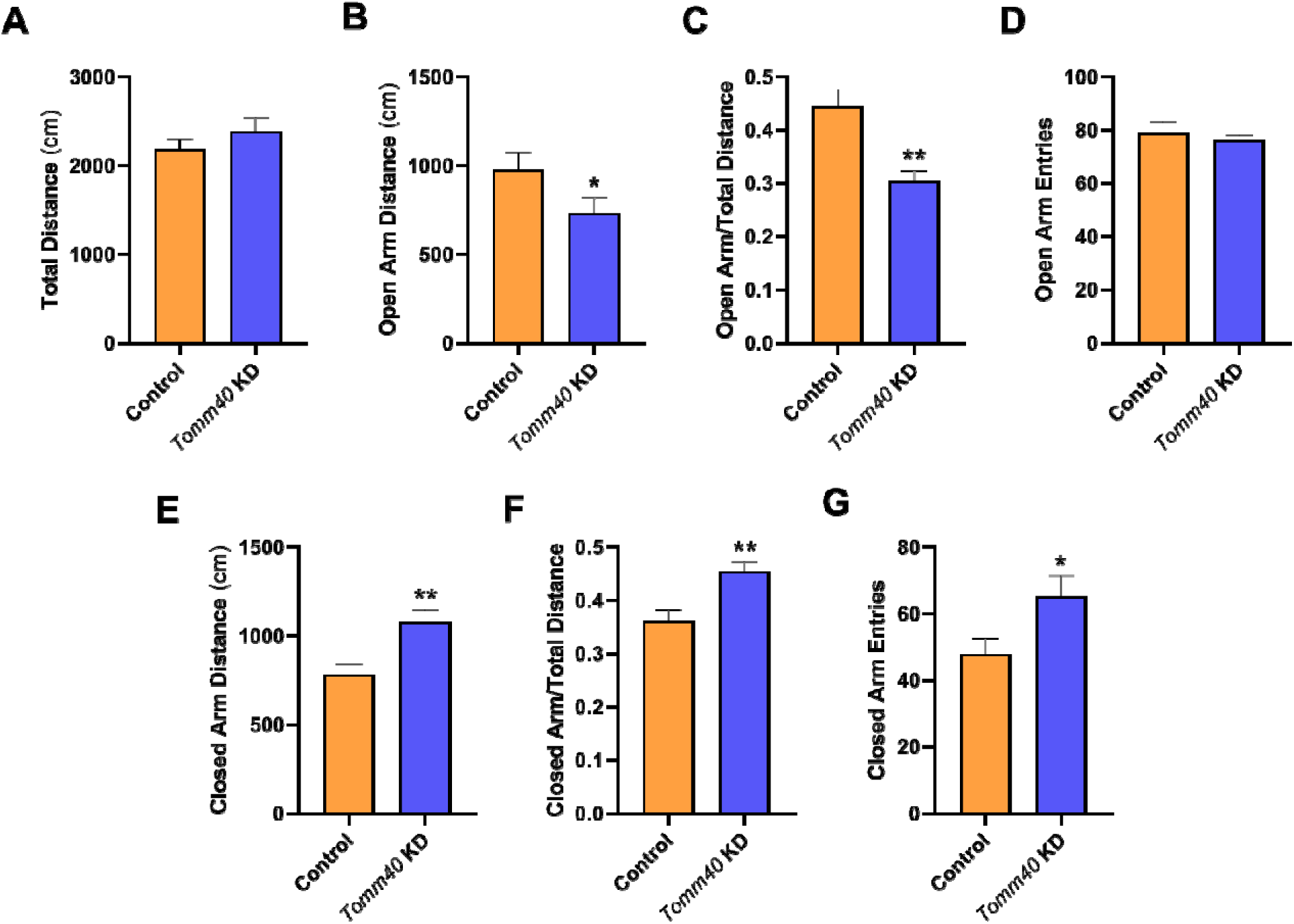
Elevated plus maze indicates anxiety-like behaviors in *Tomm40* KD mice. (A-G) EPM analysis in *Tomm40* KD vs control AAV mice: (A) Total distance (cm) travelled by mice in the EPM, (B) Total distance (cm) travelled in the open arms, (C) open arm-to-total distance ratio, (D) average number of entries into the open arms, (E) distance traveled (cm) in the closed arm, (F) closed arm to total distance ratio, and (G) number of entries into the closed arm of EPM. For all: *n=10* mice/group. *\*p<0.05, **p<0.01* vs control AVV by one-way repeated measures ANOVA, with post-hoc Student’s t-test to identify differences between groups. Data are represented as mean SEM.

#### Novel object recognition and Morris water maze tests indicate hippocampal-dependent spatial learning and declarative memory deficits in Tomm40 KD mice

We next tested learning and memory abilities in the *Tomm40* KD mice, since memory loss is one of the earliest and most distinctive hallmarks of AD^34^. During the training phase, we found that neither control nor *Tomm40* KD mice showed discrimination between 2 identical novel objects in terms of interaction time and number of bouts (**Fig. S2A-D**). During the testing phase (24 hrs after training), there was a slight, non-significant preference for the novel object in the control mice, but no differences in the number of interaction bouts between the familiar vs novel object in the *Tomm40* KD group (**Fig. 3A**). Moreover, control mice showed increased interaction time with the novel object, indicating that long-term, declarative memory was intact^35^, while *Tomm40* KD mice showed no preference for the novel object, and no difference in time spent (%) interacting with the familiar vs novel object (**Fig. 3B**). Taken together, this implies that *Tomm40* KD mice failed to discriminate between the familiar and novel object. Indeed, the novel object recognition discrimination ratio (interaction time of novel-to-familiar object) was significantly decreased in the *Tomm40* KD group compared to the controls (**Fig. 3C**), confirming declarative memory deficits in these mice. However, the *Tomm40* KD mice showed no difference from controls in the object context congruence (OCC) task (**Fig. S3**), indicating that while context discrimination is impaired, pattern separation is intact, and suggesting that Tomm40’s effect may be brain-region specific^36^. Lastly, there were no significant differences in the total number of bouts and interaction times with novel and familiar objects between the *Tomm40* KD mice and controls (**Fig. 3D & E)**.

**Figure 3.**
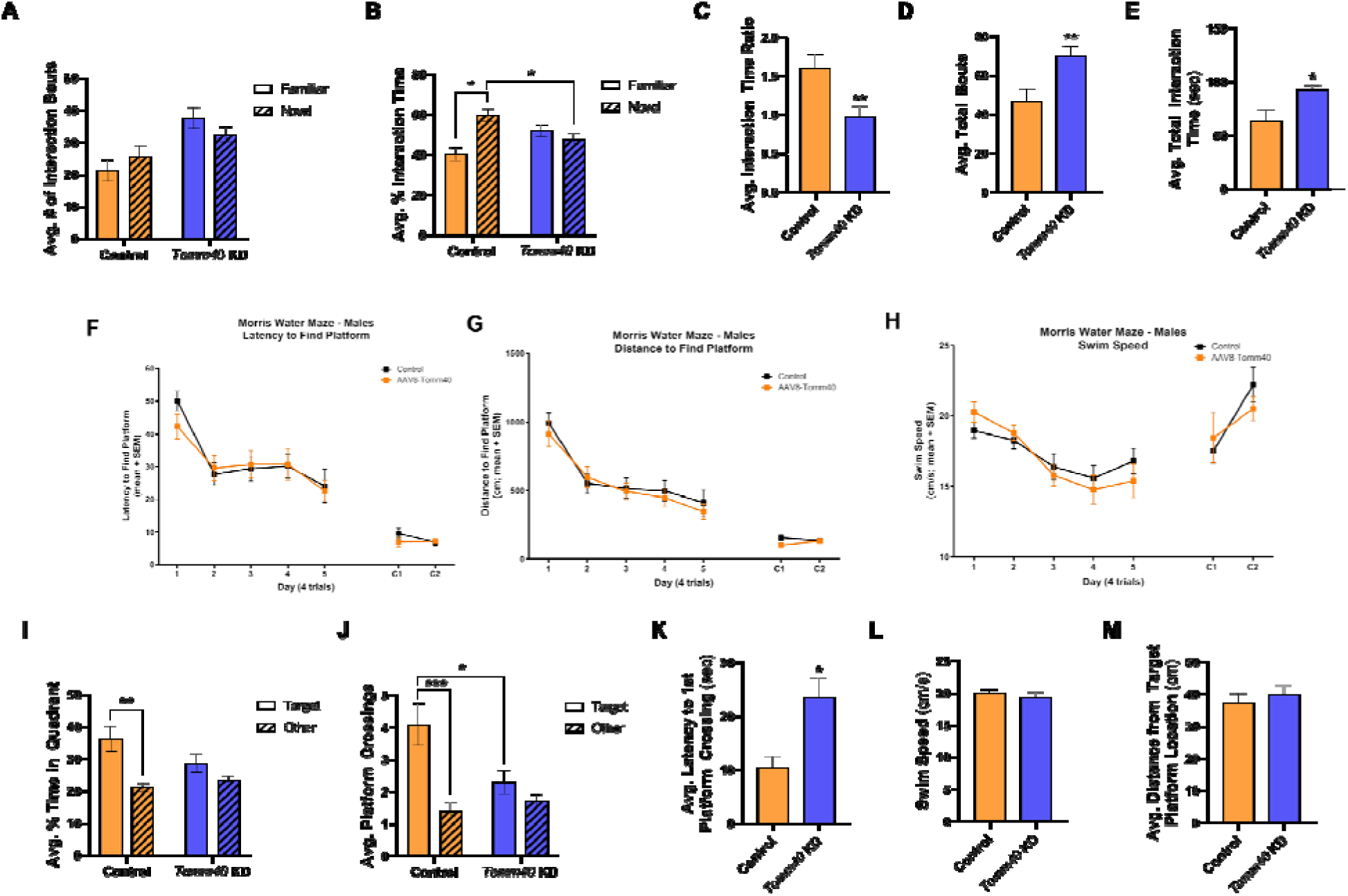
AAV8-*Tomm40* shRNA injected C57BL/6J mice display functional memory deficits and hippocampal-dependent spatial memory impairments. (A-E) NOR analysis: (A) # of interaction bouts and (B) % interaction time with familiar vs novel objects, (C) interaction time ratio to familiar (acquisition trial) and novel object (retention trial). Comparison of total bouts (D) and total interaction time (sec; E) for both familiar and novel objects between control vs. *Tomm40* shRNA AAV injected male mice. (F-H) Latency (F) and distance to find the platform (G), and swim speed (H) decreased in all groups by day 5 of training period. No differences were found betwseen both groups. Cued training days (C1 and C2) also indicated all groups learned of the platform location during acquisition training period. (I-M) MWM analysis at 24 hr: (I) % time in target vs other quadrants, (J) # of platform crossings in target vs other quadrant, (K) latency to 1^st^ platform crossing (sec), (L) swim speed throughout duration of trial, and (M) distance from target platform location (cm). For all: *n=10* mice/group. *\*p<0.05, **p<0.01, ***p<0.001* vs control AAV by one-way repeated measures ANOVA, with post-hoc Student’s t-test to identify differences between groups. Data are represented as mean SEM.

We next tested spatial learning ability and development of long-term memory in the Morris water maze (MWM)^37^, first with 5 days of training using a submerged, hidden platform and distal and proximal cues. Four trials were conducted per day in which the mice were repeatedly placed in different areas of the pool to encourage the use of allocentric (vs. egocentric) spatial learning strategies. During this training, both *Tomm40* KD and control mice showed progressively decreased latencies and distance needed to find the hidden platform over time (**Fig. 3F & G**). In addition, swimming speed did not differ between the two groups throughout the training period (**Fig. 3H**). During the acquisition training, the mice were also tested for two days to locate a cued platform and in doing so mitigate possible motor or sensory deficits affecting the performance of these mice. Latency and distance to finding the cued platform were significantly lower than in the hidden cued training (**Fig. 3F & G**) and swim speed increased (**Fig. 3H**), but there were no differences between the control and *Tomm40* KD mice. Overall, these results indicate that in the training period both *Tomm40* KD and control mice were able to reach both hidden and cued platforms and perform the MWM test successfully.

After 24 hrs from the time of acquisition training, a probe trial was conducted to assess spatial reference and long-term memory. In this test where the platform was removed, control mice spent significantly more time (%) and preference in the target quadrant (where the platform was originally) vs all other quadrants, while *Tomm40* KD mice showed no preference for any quadrant (**Fig. 3I**). Consistently, control mice showed higher number of platform crossings in the target quadrant compared to all other quadrants, while *Tomm40* KD mice showed decreased target platform crossings (**Fig. 3J**). Additionally, the time to reach the first target platform crossing was significantly longer for *Tomm40* KD than control mice (**Fig. 3K**). Swim speed and average distance from target platform location were unaffected (**Fig. 3L & M**). At the 72 and 120 hrs probe trials, control mice still showed a significant preference for target platform crossings compared to all other crossings, but *Tomm40* KD mice consistently showed no preference, confirming impaired long-term spatial memory (**Fig. S4**). Taken together, the results from the probe trials indicate significant hippocampal spatial learning and reference memory deficits^38^ in the *Tomm40* KD mice compared to controls, especially at 24 hrs post-acquisition training.

### *TOMM40* KD in human iNeurons regulates cholesterol and mitochondrial metabolism via MERCs

Using human iPSCs, we induced neural progenitor cells into fully differentiated neurons (aka iNeurons) and confirmed the protein expression of NeuN, a marker of neuronal maturation, in this cell population (**Fig. S5**). *TOMM40* KD of ∼60% in these cells was achieved using lipofectamine™ Stem transfection reagent (**Fig. 4A**), By TEM, we observed an increase in ER-mitochondria distance, and a decrease in length of MERCs and % of mitochondria with ER contact sites in *TOMM40* KD iNeurons compared to the non-target control (NTC) treated cells (**Fig. 4B-E**). Consistent with these effects, *TOMM40* KD resulted in decreased protein expression of mitofusin 1 and 2 (MFN1 and MFN2) which tethers ER to mitochondria to maintain MERCs^39^ (**Fig. 4F & G**). Since MERCs disruption promotes production of ROS^40^, we assessed cellular and mitochondria-derived ROS and found an increase in mitochondria-derived ROS with *TOMM40* KD iNeurons, with no additive effect following double KD with *MFN2*, suggesting that ROS are generated via MERCs disruption (**Fig. 4H & I**).

**Figure 4.**
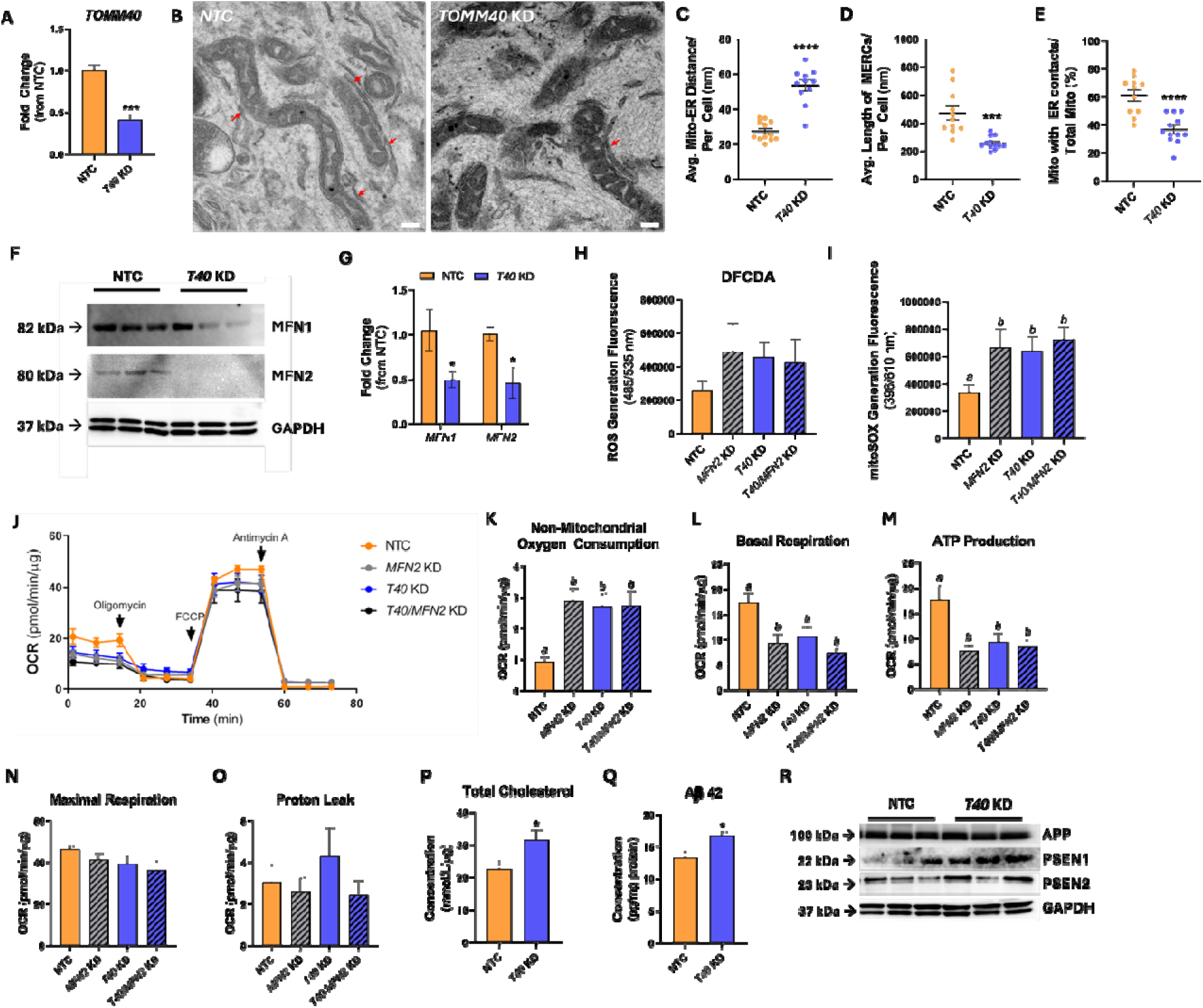
Disruption of MERCs by *TOMM40* KD impairs mitochondrial function and promotes ROS in human iPSC-derived neurons. (A) Confirmation of *TOMM40 (T40)* KD in iNeurons by ∼60%, measured by qPCR. (B) TEM micrographs of NTC and *TOMM40 (T40)* KD in iNeurons. Arrowheads indicate MERCs; scale bars, 0.5 μm. (C-E) Analysis of MERCs using ImageJ software: (C) Mito-ER distance (nm), (D) length of MERCs (nm), (E) percentage of mitochondria with ER contacts out of total mitochondria per cell. (*n = 10-12* cells/group) (F) Representative western blot of MFN1 and MFN2 protein expression in iNeurons compared to GAPDH control. (*n = 3* biological replicates) (G) mRNA transcript levels of *MFN1* and *MFN2* in iNeurons quantified by qPCR. (H) Cellular ROS and (I) mitochondria-derived ROS in *TOMM40 (T40)* and *MFN2* KD vs. NTC iNeurons was quantified by fluorescence probe, DFCDA and mitoSOX, respectively. (*n = 3* biological replicates) (J-M) Oxygen consumption rates of iNeurons transfected with *TOMM40* and/or *MFN2*, or NTC siRNAs, were quantified using the Seahorse 96e Exrtacellular Flux Analyzer. With the addition of oligomycin, FCCP, and Antimycin A + Rotenone, non-mitochondrial oxygen consumption (K), basal respiration (L), ATP production (M), maximal respiration (N), and proton leak (O), were quantified. (*n = 10-12* biological/replicates) (P) Total cholesterol quantified from iNeurons using Amplex Red Cholesterol Assay. (Q) Aβ 42 levels were quantified from iNeurons by enzyme-linked immunosorbent assay and quantified on a microplate fluorescence spectrophotometer. (R) Representative western blot of APP, PSEN1, and PSEN2 protein expression of the Aβ 42 pathway in NTC vs. *TOMM40* KD iNeurons. (*n = 3* biological replicates) For all: *\*p < 0.05, **p<0.01, ***p < 0.001, ****p < 0.0001* vs. NTC by one-way ANOVA, with post-hoc Student’s t-test to identify differences between groups. *P < 0.05* for *a* vs. *b* by two-way ANOVA, with Sidak’s multiple comparisons test. Data are represented as mean ± SEM.

Using the Seahorse Extracellular Flux Analyzer (**Fig. 4J**), we found an increase in non-mitochondrial oxygen consumption with *TOMM40* KD and/or *MFN2* double KD (**Fig. 4K**), suggesting that increased oxygen consumption may be utilized by the cell for ROS production. We also found *TOMM40* KD to impair ATP production and basal respiration in iNeurons, with no additive effect after double KD with *MFN2*, suggesting that mitochondrial dysfunction may be caused by MERCs disruption (**Fig. 4L & M**). Maximal respiration and proton leak were unaffected by *TOMM40* or *MFN2* KD in iNeurons (**Fig. 4N & O**).

Consistent with our *in vivo* findings we found an increase in intracellular cholesterol in *TOMM40* KD iNeurons (**Fig. 4P**). In addition, we found Aß 42 levels to be increased in the *TOMM40* KD vs NTC group (**Fig. 4Q**). Lastly, we showed that *TOMM40* KD promoted expression of PSEN1/PSEN2, major subunits of the y-secretase complex which cleaves amyloid precursor protein (APP) into Aß peptides, including Aß 42^41^ (**Fig. 4R**), although APP expression was not affected. In summary, these results indicate that *TOMM40* KD in iNeurons disrupts MERCs, leading to both mitochondrial dysfunction and cellular cholesterol accumulation, two causal pathways in AD pathogenesis.

### *TOMM40* KD in iNeurons increases intracellular cholesterol content via an LXR-APOE pathway

Since we previously showed that in the liver, TOMM40 regulates cholesterol pathways via LXR and its downstream transcriptional targets *APOE*, *LDLR* (cholesterol uptake), and *ABCA1* (cholesterol efflux)^42^, we first confirmed by qPCR that *TOMM40* KD upregulates the LXR pathway in iNeurons. Consistent with our findings in liver, *LXRB (NR1H2)* gene expression, but not *LXRA (NR1H3)*, was upregulated along with LXR’s downstream targets *APOE, SREBF1c, ABCA1*, and *ABCG1* (**Fig. 5A**), as well as *LDLR* and *LRP1* (**Fig. 5B**). As shown in the liver previously, *TOMM40* KD did not affect expression of *SREBF2* (sterol regulatory element-binding protein) the main transcriptional regulator of *LDLR*^43^, suggesting that upregulation of *LDLR* was instead promoted by SREBF1c^12^. Consistent with the findings in iNeurons, the LXR gene targets were also upregulated in our *Tomm40* KD mouse model (**Fig. 1J**). The role of LXR in mediating these effects in *TOMM40* KD iNeurons was confirmed by showing that downstream transcriptional targets of LXR were all downregulated following inhibition of LXR activity with 10 µM GSK 2033^44^ (**Fig. 5A**), and that the *TOMM40* KD-induced increase in cellular cholesterol content was reversed (**Fig. 5C-E**).

**Figure 5.**
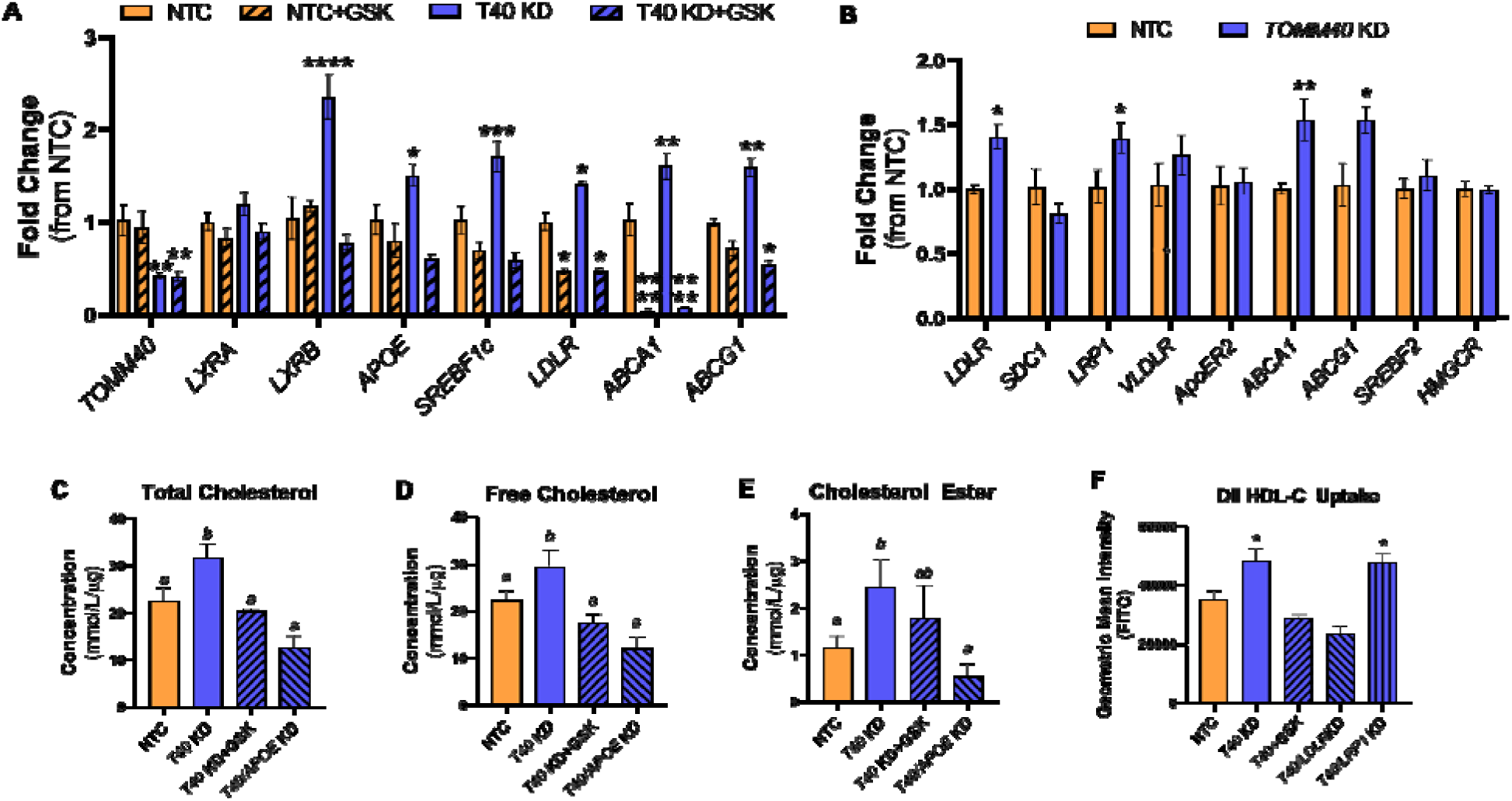
*TOMM40* KD upregulates LXRB and downstream gene targets involved in cholesterol metabolism. (A) mRNA transcript levels of *LXRA* and *LXRB* and their downstream targets in NTC vs. *TOMM40 (T40)* KD, with or without GSK2033. (B) mRNA transcripts of cholesterol uptake (*LDLR, SDC1, LRP1, VLDLR, ApoER2*), efflux (*ABCA1, ABCG1*), and synthesis (*SREBF2, HMGCR*) markers were quantified by qPCR in iNeurons treated with NTC vs. *TOMM40 (T40)* KD. (*n=3* biological replicates) (C-E) Intracellular total cholesterol (C), free cholesterol (D), and cholesterol ester (E) levels quantified from iNeurons. (*n=4* biological replicates) (F) DiI-labelled HDL-C was taken up in iNeurons and fluorescence was quantified on a microplate fluorescence spectrophotometer. (*n = 6-8* biological replicates) For all: *\*p < 0.05,**p < 0.01, *** p < 0.001, ****p < 0.0001* vs. NTC by one-way ANOVA, with post-hoc Student’s t-test to identify differences between groups. *P < 0.05* for *a* vs. *b* by two-way ANOVA, with Sidak’s multiple comparisons test. Data are represented as mean ± SEM.

To test whether *TOMM40* KD increased cellular cholesterol content via APOE-mediated effects on cholesterol uptake and/or efflux, we performed a *TOMM40/APOE* double KD in iNeurons, and found that cholesterol levels were reduced to the same extent as with the LXR antagonist, indicating that APOE plays a key role in mediating the effects of *TOMM40* KD on cellular cholesterol content (**Fig. 5C-E**). We next incubated iNeurons with DiI-labeled human HDL in order to assess effects of *TOMM40* KD on cholesterol uptake and found that cellular fluorescence increased after 4 hrs (**Fig. 5F**). This effect was reversed with double KD of *TOMM40* together with *LXR, APOE,* or *LDLR* but not *LRP1*, indicating that LXR-mediated upregulation of LDLR and its APOE ligand is responsible for increasing cholesterol uptake with *TOMM40* KD.

### *TOMM40* and *APOE4* – induced effects on cholesterol and Aß 42 levels are rescued following suppression of LDL receptor-mediated neuronal uptake

To determine whether TOMM40’s effects on neuronal cholesterol levels and AD phenotypes are dependent on the APOE isoform, we used iNeurons derived from *APOE3* (control) vs *APOE4* isogenic carriers. We found that *APOE4* iNeurons contained higher intracellular cholesterol and that the level was further increased with *TOMM40* KD (**Fig. 6A-C**). There were significant reductions of cholesterol content with *TOMM40/LDLR* double KD in both *APOE3* and *APOE4* iNeurons, but the higher level in *APOE4* cells persisted (**Fig. 6A-C**). Moreover, reductions in cholesterol content with *TOMM40/APOE* double KD were similar for the two *APOE* genotypes. Supporting these findings, we observed no differences in HDL cholesterol uptake between the APOE isoforms (**Fig. 6D**). Lastly, we found that double KD of *TOMM40* together with *APOE* or *LDLR* KD resulted in similar reductions of Aß 42 protein expression in *APOE3* and *APOE4* iNeurons (**Fig. 6E**). These results indicate that *TOMM40* KD drives up cholesterol and Aß 42 levels in iNeurons by mechanisms independent of APOE isoform, and that these effects are reversed following suppression of APOE- and LDLR-mediated cholesterol uptake.

**Figure 6.**
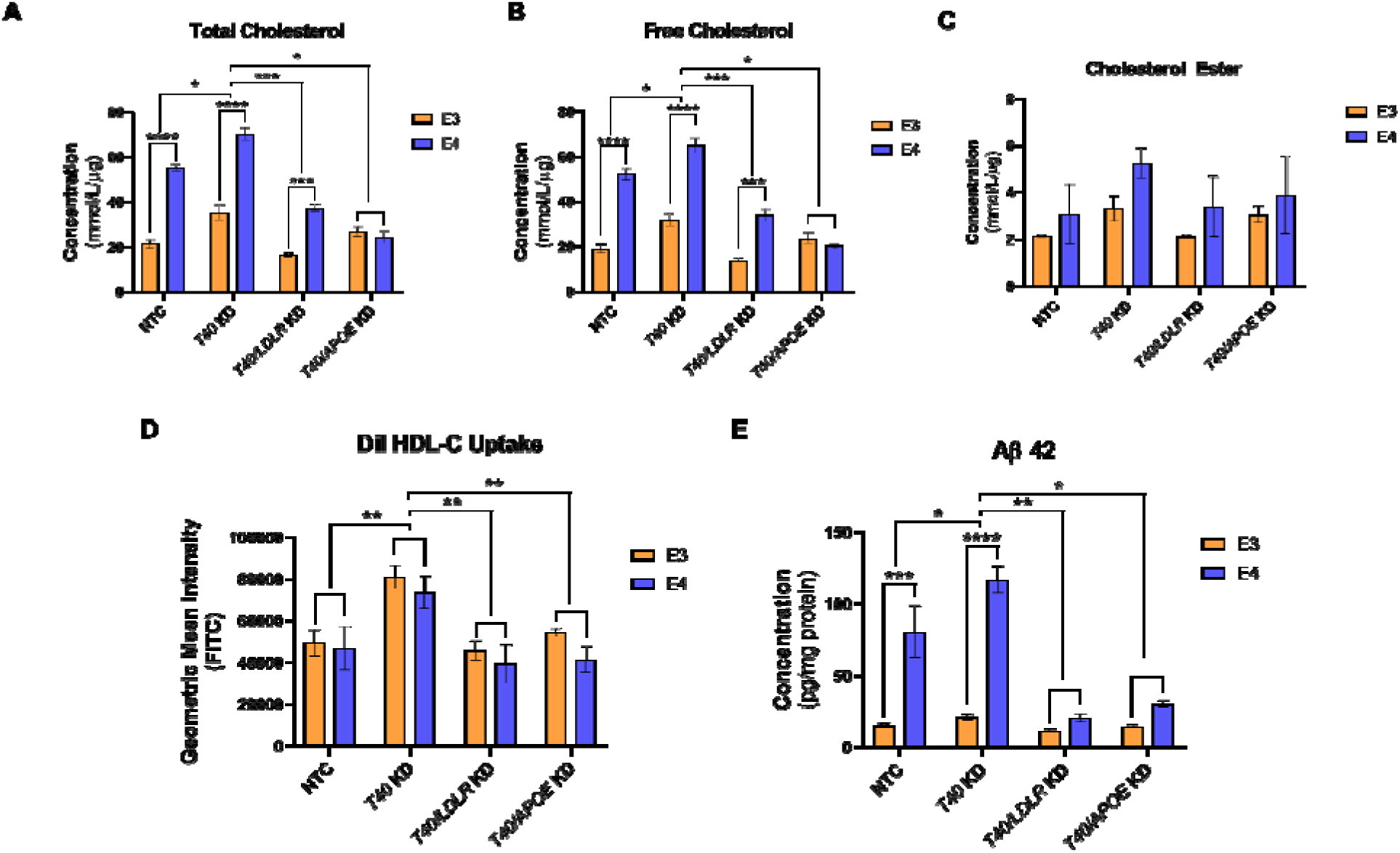
*TOMM40* KD promotes LDL uptake via an LXR-mediated pathway, mediating intracellular cholesterol and Aβ 42 levels in iNeurons. (A-C) Intracellular total cholesterol (A), free (B), and ester (C) cholesterol levels were quantified from iNeurons transfected with *NTC, TOMM40, LDLR*, and *APOE* siRNAs, singly and in combination, using Amplex Red Cholesterol Assay. (D) DiI-labelled HDL-C uptake in iNeurons transfected with *NTC, TOMM40, LDLR*, and *APOE* siRNAs, singly and in combination. (E) Aβ 42 levels were quantified from iNeurons. (*n = 6-8* biological replicates) For all: *n = 3-4* biological replicates except (B) and (F). *\*p < 0.05, **p < 0.01, ***p < 0.001, ****p < 0.0001* vs. NTC by one-way ANOVA, with post-hoc Student’s t-test to identify differences between groups. Data are represented as mean ± SEM.

### TOMM40-related phenotypes and lipid metabolism targets in a well characterized cohort from the USC ADRC stratified by clinical diagnosis (AD vs non-affected), sex, and age

To extend our *in vitro* findings to a clinical population, we utilized transcriptomic data from frozen brain tissue derived from 632 participants in a well characterized AD study cohort (ROSMAP)^45^, and found that mRNA levels of *TOMM40* were significantly correlated with the expression of the candidate cholesterol regulatory genes *LDLR*, *ABCA1, and ABCG1* (**Fig. 7A-C**), and that *TOMM40* expression was lower in brains from AD patients vs cognitively healthy controls, by bulk RNA-sequencing. These findings were evident in neurons (both excitatory and inhibitory) and not in other cell types (**Fig. 7D**). In addition, using single-nucleus RNA sequencing analysis in brains from 426 participants in ROSMAP, we found that *TOMM40* expression in inhibitory neurons was negatively associated with AD diagnosis (NIA-Reagan, Braak stage)^46,47^ and AD pathologies, including neurotic plaques, diffuse plaques, and amyloid density (**Fig. 7D**). *TOMM40* expression in excitatory neurons was strongly inversely associated with NIA-Reagan score (gold standard AD diagnosis) and amyloid density (**Fig. 7E, F**). Thus, *TOMM40* expression in excitatory neurons was found to be lower in AD participants compared to cognitively healthy controls (**Fig. 7D**) and was negatively associated with amyloid density in both cell types (**Fig. 7G, H**). Taken together, these findings provide evidence that effects of TOMM40 suppression observed in human iNeurons and mice contribute to features of AD in human brains.

**Figure 7.**
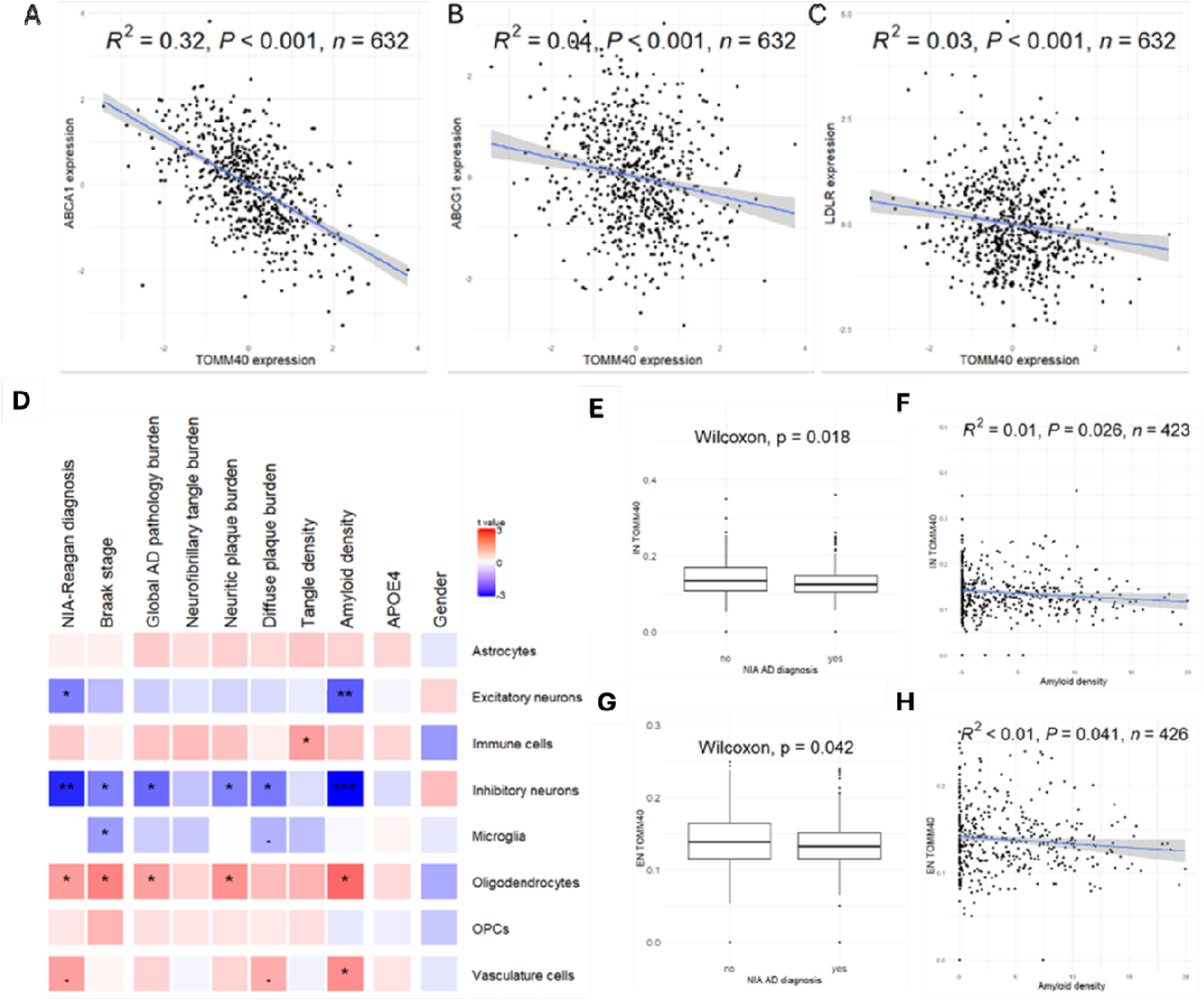
Association of *TOMM40* expression with AD factors in different brain cells. Association between the expression of TOMM40 with LXR regulate genes – ABCA1 (A), ABCG1 (B), and LDLR (C) from bulk-RNA sequencing, analyzed using a linear regression model. (D) Single-nucleus RNA sequencing analysis of the association of *TOMM40* expression levels in different brain cells with factors, including AD diagnosis, AD pathology, APOE genotype, and sex. Data shown are color coded t value (the signed effect size divided by standard error) and “fdr” adjusted p-value per each cell type (p<0.1, *p<0.05, **p<0.01, ***p<0.001). (E,G) Single-nucleus RNA sequencing analysis showing cellular *TOMM40* expression across NIA AD diagnosis in inhibitory neurons (IN, E) and excitatory neurons (EN, G). (F,H) Correlation between amyloid density and cellular *TOMM40* expression using a linear mixed effect model in IN (F), and EN (H) using single-nucleus RNA sequencing.

## DISCUSSION

Alzheimer’s disease is characterized by the accumulation of extracellular amyloid beta (Aß) plaques and neurofibrillary tangles (NFT) consisting of phosphorylated tau species. High levels of brain cholesterol have been linked to AD severity by mechanisms that result in accumulation of these Aß and tau proteins. Moreover, the strongest genetic risk factor for late onset AD (LOAD) is the *e4* allele of the gene encoding APOE which has key functions in cholesterol transport. Multiple SNPs in *APOE* associated with AD risk are in linkage disequilibrium with SNPs in *TOMM40*, which encodes a major component of the TOM complex that mediates uptake of mitochondria-targeted proteins, Moreover, at least one *TOMM40* SNP (rs 0524523) is associated with AD independent of *APOE* genotype^48^. While these genetic observations raise the possibility that TOMM40 plays a role in AD pathogenesis, the present study is the first to provide experimental evidence supporting this hypothesis.

Behavioral assessments following transient *Tomm40* whole body KD in male mice showed increased anxiety and impaired memory, features commonly observed in AD mouse models^49^. In particular, we found NOR discrimination ratio and interaction time with the novel object to be significantly lower compared to the control group, thus indicating lack of ability to discriminate novel from familiar objects and thus loss of declarative memory due to *Tomm40* suppression in the brain. However *Tomm40* KD did not impair the ability to perform the OCC task, suggesting that the dentate gyrus, which is required for discriminating pattern separation, was not affected. This is further supported by previous reports that the dentate gyrus is resistant to the formation of amyloid ß plaques and tau tangles until late stages of AD^50,51^. With the MWM test we confirmed impaired long-term spatial, reference memory which is an early marker of AD due to hippocampal atrophy^52^. Namely, the *Tomm40* KD mice showed no preference for and decreased activity in the target quadrant, indicating that the KD mice did not employ an allocentric learning strategy to solve the task. These are behavioral hallmarks of hippocampal dysfunction and potentially deficits in the perirhinal cortex which is known to coordinate visual perception and memory^53^.

In the *Tomm40* KD brain tissues there were increased levels of cholesterol and Aß 42 protein, consistent with previous evidence that increased saturation of cholesterol ester in the plasma membrane of neural cells promotes cleavage of transmembrane amyloid precursor protein (APP) by γ-secretase activity^54^ generating Aß 40/42 products. Aß 42, which is two hydrophobic amino acids longer than Aß 40 and is considered the major neurotoxic amyloid peptide that promotes neurodegeneration in AD^55^. Unlike Aß 42, no differences were observed in p-tau levels, possibly due to insufficient duration of *Tomm*40 KD^56^. Contrary to Aß 42 aggregates that can be detected early in AD pathogenesis, p-tau levels are detected in later stages of AD, since the amyloid-ß pathway affects tau phosphorylation downstream via the CDK5 phosphorylation pathway^57,58^. This has been confirmed in a clinical prospective study that measured cerebrospinal fluid levels in patients with mild cognitive impairment over 10 years, showing Aß 42 but not p-tau species to be significantly increased in the first 5-10 years prior to onset of AD^59^. Collectively, the characterization of major AD phenotypes in our *Tomm40* KD mouse model supports a potential causal role for impaired TOMM40 levels and/or functionality in AD.

As a model system relevant to humans, we next employed human iPSC-derived iNeurons to investigate the cellular and molecular mechanisms for the causal effects of *TOMM40* suppression observed *in vivo*. Likewise, *TOMM40* suppression promoted cholesterol uptake in iNeurons which we found to be due to upregulation of *LDLR* and its *APOE* ligand. This was confirmed by finding reduced cholesterol content following suppression of *LDLR-* or *APOE-* mediated HDL cholesterol uptake in *TOMM40* KD iNeurons. While *TOMM40* KD also increased expression of *LRP1,* another gene encoding APOE-binding protein that facilitates cholesterol uptake^60^, KD of this gene had no effect on HDL uptake or cholesterol content. Together with increased cholesterol uptake, higher Aß 42 content was found in *TOMM40* KD iNeurons, and this was restored to baseline level following *LDLR-* or *APOE-* suppression, suggesting that Aß 42 content is regulated by intracellular cholesterol in iNeurons.

We have shown previously that hepatic *TOMM40* KD upregulates *LDLR* and *APOE* transcription via LXR, a nuclear receptor that regulates cellular cholesterol metabolism^12^. Downstream transcriptional targets of LXR include *APOE* and *SREBF1c*, which has been shown, along with *SREBF2*, to promote the transcription of *LDLR* in human hepatoblastoma cells and hepatocytes^12,61^. We here have demonstrated that this pathway is conserved in iNeurons, and confirmed that as in liver, LXRB is the subtype specifically responsible for upregulating *APOE* and *LDLR* gene expression. Expression levels of *ABCA1* and *ABCG1*, LXR transcriptional targets that are key regulators of cellular cholesterol efflux were also upregulated by *TOMM40* KD, raising the possibility that this effect could have attenuated the cholesterol accumulation in iNeurons. However, it has been shown that ABCA1-mediated cholesterol efflux in neurons is suppressed due to increased ABCA1 endolysosomal trafficking that results from increased oxysterol activation of the MTORC1 cellular senescence pathway^62^. Thus, our study has identified a novel molecular pathway whereby *TOMM40* KD promotes greater neural cell cholesterol content via increased uptake due to upregulation of both LDLR and its APOE-ligand, though further studies will be needed to determine the extent to which this effect might be offset by ABCA1- and/or ABCG1-mediated cholesterol efflux.

Also consistent with our previous findings in hepatocytes^12^, we demonstrated that *TOMM40* KD in iNeurons increased cellular cholesterol content by disrupting MERCs, and that MERCs disruption resulted in increased generation of cellular and mitochondria-derived ROS that drive production, by cholesterol oxidation, of oxysterols, potent activators of LXR^63,64^. Importantly we found that disruption of MERCs by either *MFN2*^65^ or *TOMM40* KD results in impaired mitochondrial function, as manifest by reduced oxidative phosphorylation and mitochondrial respiration. Mitochondrial dysfunction has been implicated in AD pathogenesis^66–68^ and has been shown to accelerate the accumulation of Aß plaques and tau tangles by disrupting protein degradation pathways^69,70^.

Given the strong association of the APOE4 isoform with AD risk, we next used iNeurons derived from carriers of the *APOE4* allele and *APOE3* homozygotes to test for possible differences in cholesterol content and interactions with *TOMM40* expression. We found that cholesterol levels were higher in the *APOE4* iNeurons, and remained so after *TOMM40* KD. Moreover, KD of *LDLR* or *APOE* reduced Aß 42 levels to the same extent in both cell types. These results indicate that *TOMM40* KD increases cholesterol and Aß 42 levels in iNeurons by mechanisms independent of and additive to the presence of the *APOE4* isoform.

Taken together, our findings in iNeurons support a novel mechanism whereby disruption of MERCs by suppression of *TOMM40* expression leads to effects associated with AD pathogenesis, including mitochondrial dysfunction and LXRB-mediated increases in cholesterol and Aß 42 content that are independent of the APOE isoform. Moreover these in *vitro* effects are consistent with the adverse impact of *Tomm40* KD on measures of cognition *in vivo* in mice. We therefore sought to assess the clinical relevance of these findings by studies of human brain samples. Notably we found that in both inhibitory and excitatory neurons there was a strong inverse association of *TOMM40* expression with AD diagnosis and with increased cellular content of neurotic plaques, diffuse plaques, and amyloid density, all phenotypes of AD pathology. While the magnitude of these associations was not great, they may yet be sufficient to have significant clinical impact over time. Finally, utilizing RNA-seq data from a bank of human brain tissues, we confirmed significant negative correlations between *TOMM40* mRNA levels and expression of cholesterol regulatory genes including *LDLR, ABCA1*, and *ABCG1.* Collectively, these clinical findings further support our *in vitro* experiments that have shown TOMM40 to be a mediator of neuronal cholesterol metabolism and the amyloid pathway.

A strength of this work is the use of studies in mice, human iNeurons, and human brain tissue that provide complementary information supporting a novel role for reduced neuronal *TOMM40* expression in the pathogenesis of AD. The study does however have several limitations. While neurons are heavily affected in AD and are the main sites for neurodegeneration^71^, it will be necessary to also determine the effects of *TOMM*40 KD in microglia and astrocytes to provide a more comprehensive assessment of TOMM40’s role in AD pathogenesis. In addition, since the *in vivo* studies were limited to transient *TOMM40* KD in male mice, future studies will be needed to test for possible sex differences in response, and to assess longer term impact on AD pathology using stable KD or knock-out mouse models. Finally, a larger number of human brain samples will be needed to provide sufficient power for testing the impact of *TOMM40* genotypes that have been associated with AD risk.

## CONCLUSIONS

This study provides evidence that reduced *TOMM40* expression in the brain directly suppresses traits associated with AD pathogenesis via effects on cholesterol metabolism and mitochondrial function – two interconnected biological pathways that are increasingly implicated in AD risk. Moreover, the higher cellular content of cholesterol and Aß 42 resulting from *TOMM40* downregulation are independent of, and additive to, the effects of the APOE4 isoform. Finally, the findings suggest that maintenance of MERCs plays a pivotal role in limiting neuronal cholesterol uptake and oxidative damage, thereby suppressing progression of AD.

## AUTHOR CONTRIBUTIONS

N.V.Y. and R.M.K. conceived the idea. N.V.Y designed, performed, and analyzed the experiments, and wrote the original manuscript draft. S.W. and B.L. performed, analyzed, and interpreted RNA-seq experiments. J.S. and L.D. performed and analyzed the behavioral animal studies. J.H.O. and A.H. assisted with iPSC experiments and manuscript writing. Y.H. assisted in experimental design, data interpretation, and editing of manuscript. R.M.K. carried out project supervision, data interpretation, manuscript editing and funding acquisition.

## ACKNOWLEDGEMENTS

We thank Reena Zalpuri and Danielle Jorgens at the University of California, Berkeley, Electron Microscopy Laboratory for their advice and assistance in electron microscopy sample preparation and data collection. Justin Kim, Brandon Kamada, and Timothy Jang assisted with sample preparation, lipid extractions, cholesterol quantification, and ELISAs. Dr. Marisa Medina kindly provided use of her laboratory equipment and space.

## CONSENT STATEMENT

No human subjects were involved in this study.

## DATA AVAILABILITY STATEMENT

Data will be provided upon reasonable request to the corresponding authors.

