## Supplemental Figures for "*TOMM40* suppression promotes neuronal cholesterol imbalance and molecular and behavioral phenotypes of Alzheimer’s disease"

### Table S1. RT-qPCR primers employed in this study.

| Oligonucleotides | Primer Sequence (5' to 3') | Oligonucleotides | Primer Sequence (5' to 3') |
|---|---|---|---|
| **Human:** | | **Mouse:** | |
| *TOMM40 (forward)* | AGG AGG GCA CTG TCA TGT CT | *Tomm40 (forward)* | GAA GAT GGG AGC TGC GGA T |
| *TOMM40 (reverse)* | TGG TCA CTG GCT TTG TGG TA | *Tomm40 (reverse)* | AAG TGG TAG TTG GAC TCC CC |
| *MFN1 (forward)* | TGG CTA AGA AGG CGA TTA CTG C | *Ldlr (forward)* | TTG TGT GTG ATG GAG ACC GA |
| *MFN1 (reverse)* | TCT CCG AGA TAG CAC CTC ACC | *Ldlr (reverse)* | CGT CAA CAC AGT CGA CAT CC |
| *MFN2 (forward)* | GCA CTT TGT CAC TGC CAA GA | *Lxra (forward)* | GAA ATG CCA GGA GTG TCG AC |
| *MFN2 (reverse)* | CAC TTT CAT GTG CCT CCG AG | *Lxra (reverse)* | CAC CAG CTT CTC GAT CAT GC |
| *LXRA (forward)* | TGA GAG TAT CAC CTT CCT CA | *Lxrb (forward)* | GGC TTG CAG GTG GAA TTC AT |
| *LXRA (reverse)* | AGA AGA TGC TGA TAG CAA TG | *Lxrb (reverse)* | ATA CTC TGC ATC GTC CAG GC |
| *LXRB (forward)* | AAC TAA TGA TCC AGC AGT TG | *Apoe (forward)* | GAT CAG CTC GAG TGG CAA AG |
| *LXRB (reverse)* | ATC TCC TGG ACT GAG ATG AT | *Apoe (reverse)* | TTG TGT GAC TTG GGA GCT CT |
| *APOE (forward)* | CTC AGC TCC CAG GTC ACC | *Abca1 (forward)* | GGG AAG AGA GCA TGT GGA GT |
| *APOE (reverse)* | GGG TCA GTT GTT CCT CCA GT | *Abca1 (reverse)* | GTT GCC GCC ACT GTA GTT AC |
| *ABCA1 (forward)* | TGT AAT GCC AAC AAC CCC TG | *18s (forward)* | AGT CCC TGC CCT TTG TAC ACA |
| *ABCA1 (reverse)* | TGT CCT TCA TGC TGG TGT CT | *18s (reverse)* | CGA TCC GAG GGC CTC ACT A |
| *ABCG1 (forward)* | GGT TCT TCG TCA GCT TCG AC | | |
| *ABCG1 (reverse)* | GTT TCC TGG CAT TCA GGT GT | | |
| *SREBF1c (forward)* | ACA CAG CAA CCA GAA ACT CAA G | | |
| *SREBF1c (reverse)* | AGT GTG TCC TCC ACC TCA GTC T | | |
| *LDLR (forward)* | TCT CTT CCA CAA CCT CAC CC | | |
| *LDLR (reverse)* | CAG GTA AAC TTG GGC GAG TG | | |
| *SREBF2 (forward)* | TTT CTC CCA CCT CAG TTC CC | | |
| *SREBF2 (reverse)* | TTG GAC TTG AGG CTG AAG GA | | |
| *HMGCR (forward)* | AGC CAT TTT GCC CGA GTT TT | | |
| *HMGCR (reverse)* | GCG ACT GTG AGC ATG AAC AA | | |
| *SDC1 (forward)* | GCT ATT CCC ACG TCT CCA GA | | |
| *SDC1 (reverse)* | CCA CTT CTG GCA GGA CTA CA | | |
| *LRP1 (forward)* | CTG TAT CTC AAA GGG CTG GC | | |
| *LRP1 (reverse)* | AAC ACA CAG CTC AGT ACC CA | | |
| *VLDLR (forward)* | TGT GAA CCC TCC CAA TTC CA | | |
| *VLDLR (reverse)* | CTG CCG TCA ACA CAG TCT TC | | |
| *ApoER2 (forward)* | ATG GGC CTC CCC GAG CC | | |
| *ApoER2 (reverse)* | TCA GGG TAG TCC ATC ATC TTC AAG GC | | |
| *GAPDH (forward)* | GTG GTC TCC TCT GAC TTC AAC A | | |
| *GAPDH (reverse)* | CTC TTC CTC TTG TGC TCT TGC T | | |

**Table S2. Antibodies employed in this study for immunohistochemistry.**

| REAGENT or RESOURCE | SOURCE | IDENTIFIER |
|---|---|---|
| **Antibodies** | | |
| Anti-APP | Cell Signaling Technology | 2452 |
| Anti-GAPDH | Santa Cruz | sc32233 |
| Anti-MFN1 | Cell Signaling Technology | 14739 |
| Anti-MFN2 | Cell Signaling Technology | 9482 |
| Anti-NeuN | Cell Signaling Technology | 94403 |
| Anti-PSEN1 | Cell Signaling Technology | 5643 |
| Anti-PSEN2 | Cell Signaling Technology | 9979 |
| Anti-rabbit IgG-HRP | Cell Signaling Technology | 7074 |
| Anti-mouse IgG-HRP | Cell Signaling Technology | 7076 |

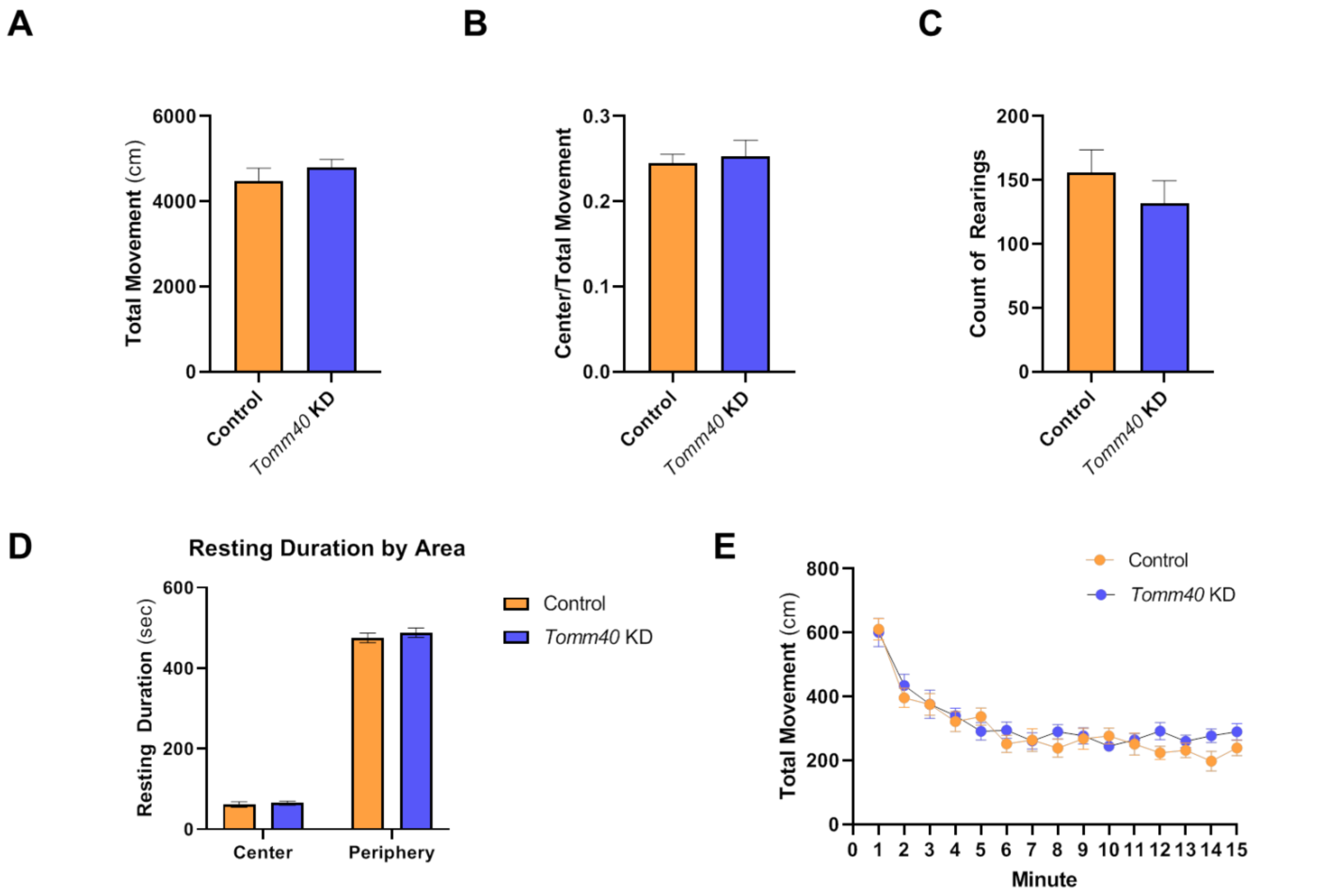


**Figure S1. Open field test analysis of *Tomm40* KD male mice.** (A) total movement (cm), (B) center-to-total movement ratio, (C) count of rearings, (D) resting duration (cm), and (E) total movement (cm) by both groups of mice were assessed. For all: *n=10* mice/group. *Tomm40* KD vs control AAV by one-way repeated measures ANOVA, with post-hoc Student's t-test to identify differences between groups. Data are represented as mean ± SEM.

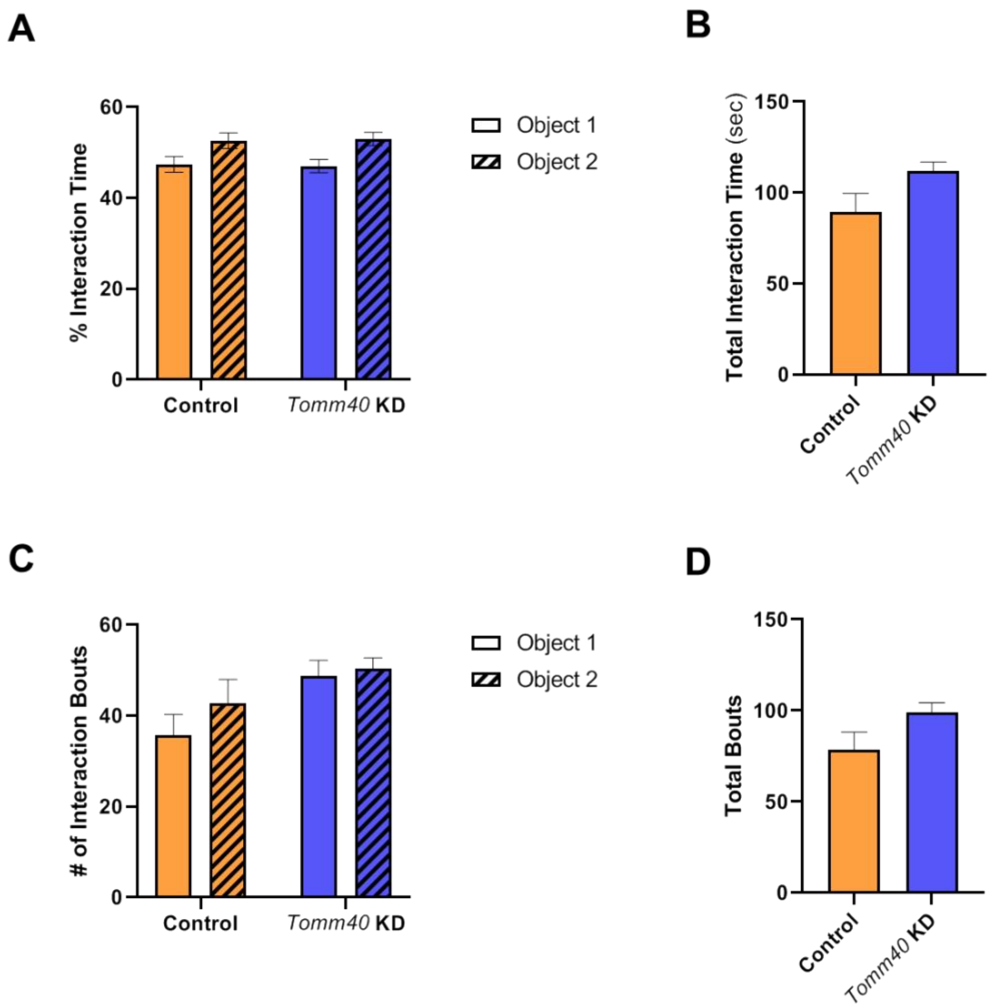


**Figure S2. No preferences in objects indicated during ORT training.** Analysis indicate both control and *Tomm40* KD male mice spend equal time (A) and # of interaction bouts (C) with both object 1 and 2 during ORT training period, 24 hrs prior to testing. Total interaction time (sec; B) and total bouts (D) by both groups showed no differences in learning or preference for both objects. For all: *n=10* mice/group. *Tomm40* KD vs control AAV by one-way repeated measures ANOVA, with post-hoc Student's t-test to identify differences between groups. Data are represented as mean ± SEM.

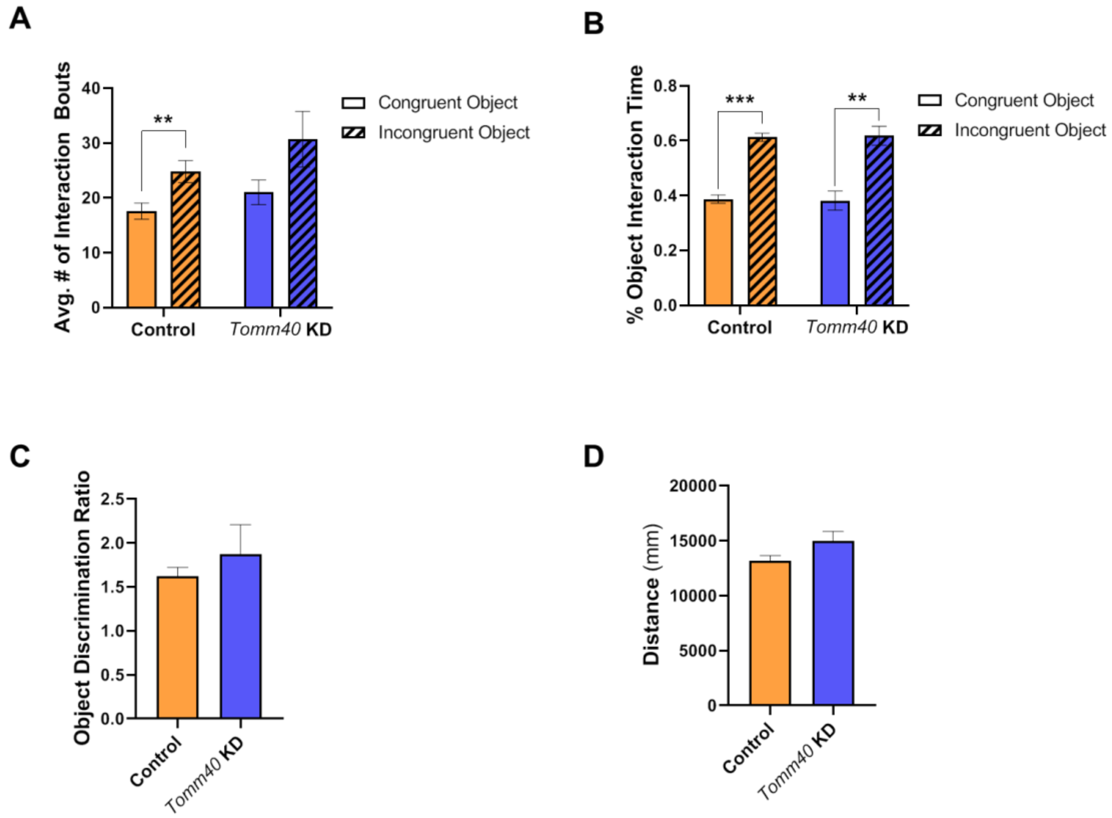


**Figure S3. No differences in performing object context congruence tasks in *Tomm40* KD mice.** Analysis indicate both control and *Tomm40* KD male mice have equal # of interaction bouts (A) and % interaction time (B) with congruent vs. incongruent objects. No differences in pattern separation or context discrimination when object discrimination ratio (C) and total distance travelled during the test (mm; D) were assessed. For all: $n=10$ mice/group. *Tomm40* KD vs control AAV by one-way repeated measures ANOVA, with post-hoc Student's t-test to identify differences between groups. Data are represented as mean $\pm$ SEM.

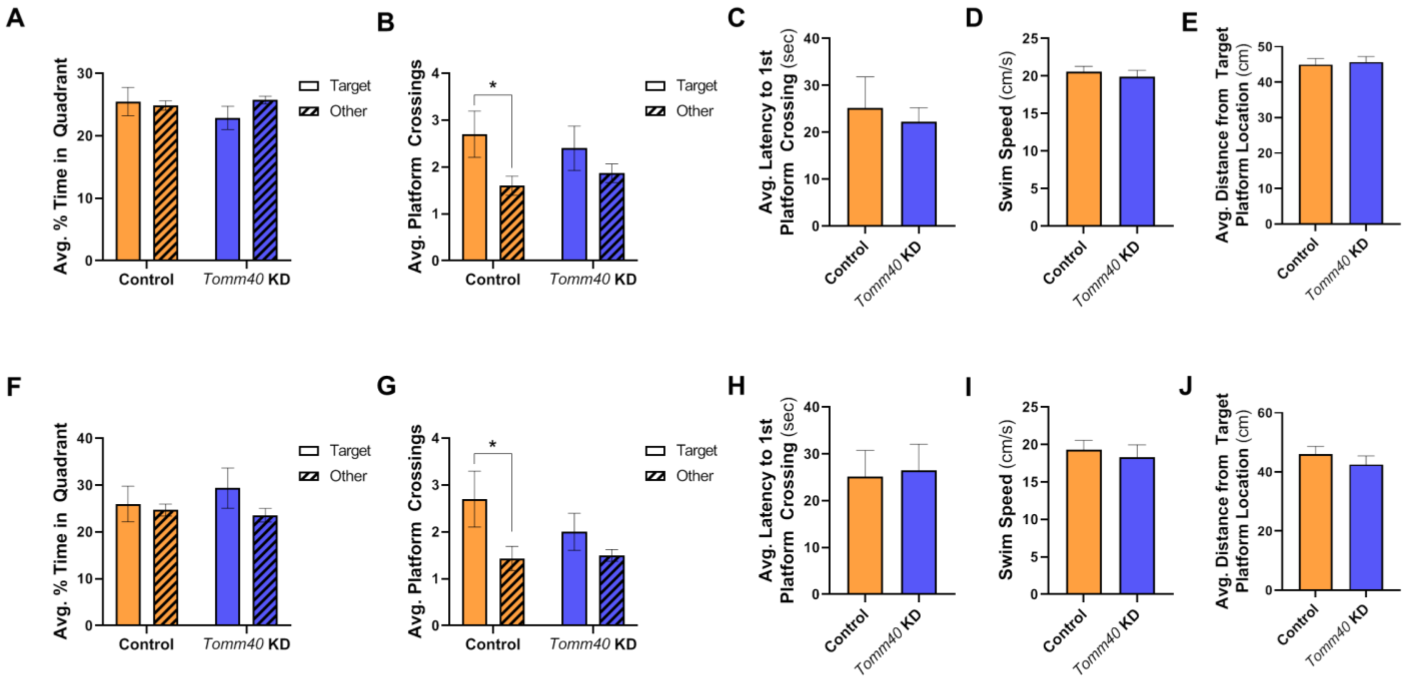


**Figure S4. Results of MWM analysis at 72 and 120 hrs probe trials.** Analysis at 72 hrs probe trial: time in quadrant (%; A), number of platform crossings (B), latency to 1$^{st}$ platform crossing (cm; C), swim speed (cm/s; D), and distance from target platform location (cm; E) were averaged across all mice in each group. Analysis at 120 hrs probe trial: time in quadrant (%; F), number of platform crossings (G), latency to 1$^{st}$ platform crossing (cm; H), swim speed (cm/s; I), and distance from target platform location (cm; J) were averaged across all mice in each group. For all: *n=10* mice/group. **p<0.05* vs control AAV by one-way repeated measures ANOVA, with post-hoc Student's t-test to identify differences between groups. Data are represented as mean ± SEM.

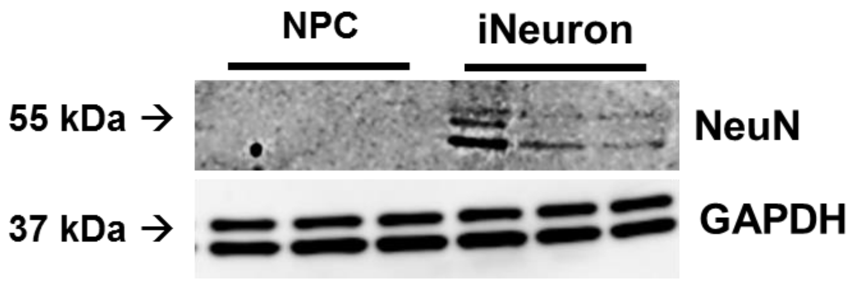


**Figure S5. Representative western blot of NeuN expression confirming differentiation of mature iNeurons from neural progenitor cells (NPC), compared to GAPDH control.**
